# Adjuvanting a subunit SARS-CoV-2 nanoparticle vaccine to induce protective immunity in non-human primates

**DOI:** 10.1101/2021.02.10.430696

**Authors:** Prabhu S. Arunachalam, Alexandra C. Walls, Nadia Golden, Caroline Atyeo, Stephanie Fischinger, Chunfeng Li, Pyone Aye, Mary Jane Navarro, Lilin Lai, Venkata Viswanadh Edara, Katharina Röltgen, Kenneth Rogers, Lisa Shirreff, Douglas E Ferrell, Samuel Wrenn, Deleah Pettie, John C. Kraft, Marcos C. Miranda, Elizabeth Kepl, Claire Sydeman, Natalie Brunette, Michael Murphy, Brooke Fiala, Lauren Carter, Alexander G White, Meera Trisal, Ching-Lin Hsieh, Kasi Russell-Lodrigue, Christopher Monjure, Jason Dufour, Lara Doyle-Meyer, Rudolph B. Bohm, Nicholas J. Maness, Chad Roy, Jessica A. Plante, Kenneth S. Plante, Alex Zhu, Matthew J. Gorman, Sally Shin, Xiaoying Shen, Jane Fontenot, Shakti Gupta, Derek T. O’Hagan, Robbert Van Der Most, Rino Rappuoli, Robert L. Coffman, David Novack, Jason S. McLellan, Shankar Subramaniam, David Montefiori, Scott D. Boyd, JoAnne L. Flynn, Galit Alter, Francois Villinger, Harry Kleanthous, Jay Rappaport, Mehul Suthar, Neil P. King, David Veesler, Bali Pulendran

## Abstract

The development of a portfolio of SARS-CoV-2 vaccines to vaccinate the global population remains an urgent public health imperative. Here, we demonstrate the capacity of a subunit vaccine under clinical development, comprising the SARS-CoV-2 Spike protein receptor binding domain displayed on a two-component protein nanoparticle (RBD-NP), to stimulate robust and durable neutralizing antibody (nAb) responses and protection against SARS-CoV-2 in non-human primates. We evaluated five different adjuvants combined with RBD-NP including Essai O/W 1849101, a squalene-in-water emulsion; AS03, an alpha-tocopherol-containing squalene-based oil-in-water emulsion used in pandemic influenza vaccines; AS37, a TLR-7 agonist adsorbed to Alum; CpG 1018-Alum (CpG-Alum), a TLR-9 agonist formulated in Alum; or Alum, the most widely used adjuvant. All five adjuvants induced substantial nAb and CD4 T cell responses after two consecutive immunizations. Durable nAb responses were evaluated for RBD-NP/AS03 immunization and the live-virus nAb response was durably maintained up to 154 days post-vaccination. AS03, CpG-Alum, AS37 and Alum groups conferred significant protection against SARS-CoV-2 infection in the pharynges, nares and in the bronchoalveolar lavage. The nAb titers were highly correlated with protection against infection. Furthermore, RBD-NP when used in conjunction with AS03 was as potent as the prefusion stabilized Spike immunogen, HexaPro. Taken together, these data highlight the efficacy of the RBD-NP formulated with clinically relevant adjuvants in promoting robust immunity against SARS-CoV-2 in non-human primates.

---

Subunit vaccines are amongst the safest and most widely used vaccines ever developed. They have been highly effective against a multitude of infectious diseases such as Hepatitis-B, Diphtheria, Pertussis, Tetanus and Shingles in diverse age groups, from the very young to the very old. An essential component of subunit vaccines is the adjuvant, an immune-stimulatory agent which enhances the magnitude, quality and durability of the immune responses induced by vaccination even with lower doses of antigen^1^. Therefore, the development of a safe and effective subunit vaccine against SARS-CoV-2 would represent an important step in controlling the COVID-19 pandemic. The most widely used adjuvant, Aluminium salts (Alum), has been used in billions of doses of vaccines over the last century. During the past decade, several novel adjuvants have been developed including the α-tocopherol containing squalene-based oil-in-water adjuvant AS03^2^, and the toll-like receptor (TLR)-9 ligand CpG 1018^3,4^, which are included in licensed vaccines against pandemic influenza and Hepatitis-B, respectively. In particular, both AS03 and CpG 1018 are currently being developed as adjuvants for use in candidate subunit SARS-CoV-2 vaccines; however, their capacity to stimulate protective immunity against SARS-CoV-2 remains unknown.

We recently described SARS-CoV-2 RBD-16GS-I53-50 (RBD-NP), a subunit vaccine in which 60 copies of the SARS-CoV-2 RBD are displayed in a highly immunogenic array using a computationally designed self-assembling protein nanoparticle (hereafter designated RBD-NP)^5^. Pre-clinical evaluation in mice showed that the vaccine elicits 10-fold higher nAb titers than the two-proline (2P) prefusion-stabilized spike (which is used by most vaccines being developed) at a 5-fold lower dose and protects mice against mouse-adapted SARS-CoV-2 challenge^5^. In the current study, we evaluated the capacity of AS03, CpG 1018 formulated in Alum, as well as the squalene-in-water emulsion (O/W), the TLR-7 agonist adsorbed to Alum (AS37)^6^ and Alum to promote protective immunity against SARS-CoV-2 in non-human primates (NHPs).

## Robust antibody responses to RBD-NP formulated with different adjuvants

To assess the immunogenicity and protective efficacy of RBD-NP vaccination with different adjuvants, we immunized 29 male *Rhesus macaques* (RMs) with 25 µg RBD antigen (71 µg of total RBD-NP immunogen; Extended Data Fig. 1) formulated with one of the following five adjuvants: O/W, AS03, AS37, CpG 1018-Alum (CpG-Alum) or Alum (Fig. 1a). Four additional animals were administered with saline as a control. All the immunizations were administered via intramuscular route on days 0 and 21 in forelimbs. Four weeks after the booster immunization, we challenged the animals with SARS-CoV-2 via intratracheal/intranasal (IT/IN) routes. Five of the ten animals immunized with AS03-adjuvanted RBD-NP were not challenged to allow evaluation of the durability of the vaccine-elicited immune responses and will be challenged at a distal time point.

**Fig. 1.**
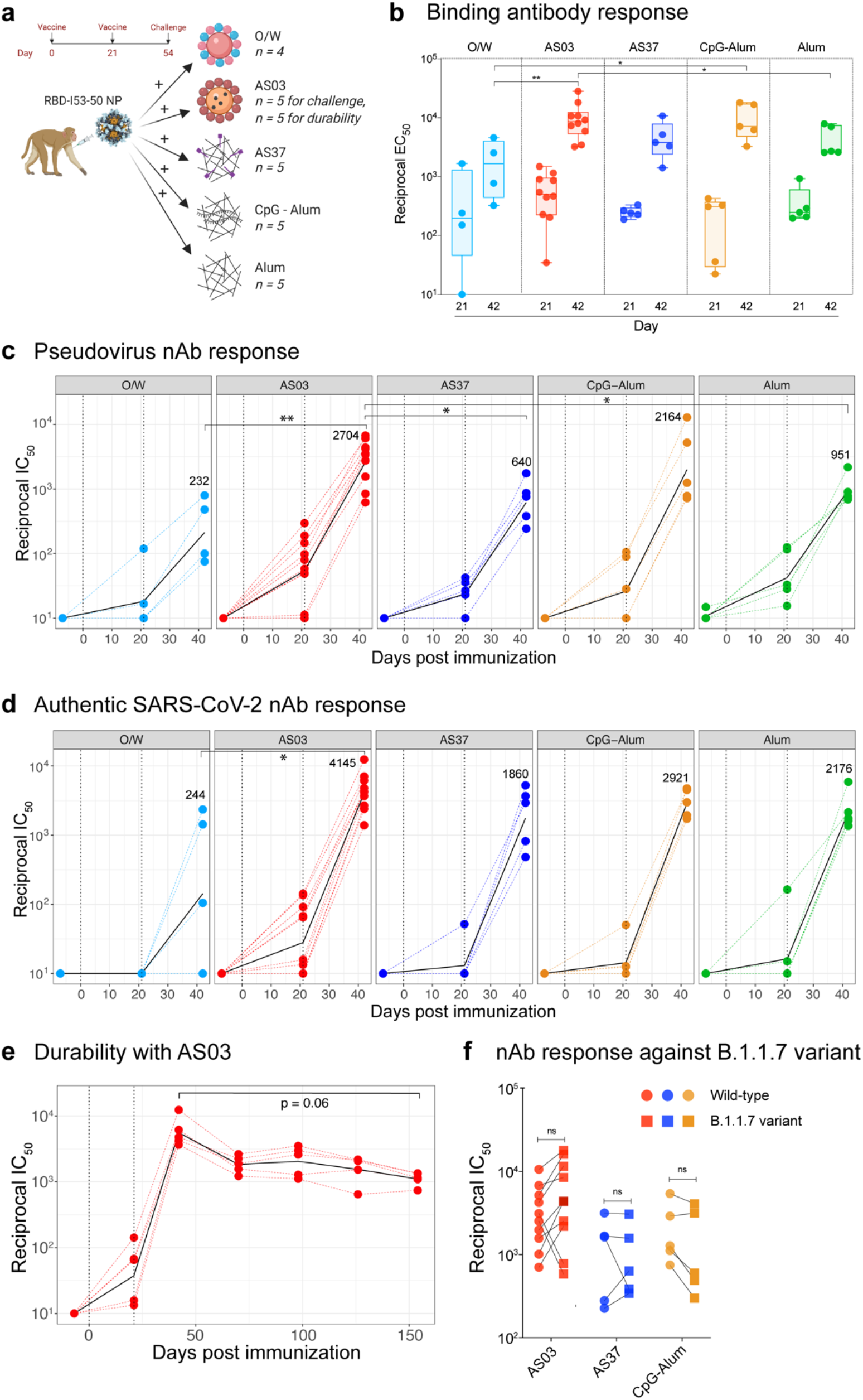
SARS-CoV-2 RBD-NP immunization induces robust antibody responses. **a**, Schematic representation of the study design. **b**, SARS-CoV-2 S-specific IgG titers (plotted as reciprocal EC_50_) in sera collected at days 21 and 42 measured by ELISA. The box shows median and 25^th^ and 75^th^ percentiles and the error bars show the range. **c - d**, Serum nAb titers (plotted as reciprocal IC_50_) determined using a SARS-CoV-2 S pseudovirus (**c**) and authentic SARS-CoV-2 (**d**) entry assay at day −7, 21 and 42. In **c** and **d**, the black line represents the geometric mean of all data points. The numbers represent geometric mean titers on day 42. Asterisks represent the statistically significant differences between two groups analyzed by two-sided Mann-Whitney rank-sum test (* p < 0.05, ** p < 0.01). **e**, SARS-CoV-2 S-specific nAb titers against authentic SARS-CoV-2 virus measured at time points indicated on X-axis. Statistical difference between the time points was analyzed by two-sided Wilcoxon matched-pairs signed-rank. **f**, Serum nAb titers against the wild-type (circles) or the B1.1.7 (squares) variant live-virus measured in serum collected at day 42, 3 weeks following secondary immunization. The statistical differences between wild-type and variant within each group were analyzed by two-sided Wilcoxon matched-pairs signed-rank test (* p < 0.05).

Evaluation of binding antibody responses to vaccination showed that S-specific IgG was detected 21 days after primary immunization in all vaccination groups and increased in magnitude after boosting (Fig. 1b). AS03 induced the highest magnitude of binding IgG (GMT EC_50_ 1:8551) on day 42, and O/W induced the lowest (GMT EC_50_ 1:1308) response. Binding antibodies in the AS37, CpG-Alum, and Alum groups were comparable to AS03 in magnitude. In addition to S-specific IgG, we also measured antibody response to the I53-50 protein nanoparticle (NP) scaffold. Anti-NP antibody titers were elicited in all the groups albeit at a lower magnitude (1.7-fold lower on average) in comparison to the anti-Spike antibody titers among the different adjuvant groups at day 42 (Extended Data Fig. 2a). The anti-NP antibody response correlated strongly with S-specific binding antibody responses (Extended Data Fig. 2b).

RBD-NP immunization induced detectable nAb responses against a SARS-CoV-2 S pseudotyped virus^7^ after primary immunization, which significantly increased in all groups after the booster immunization (Fig. 1c). In particular, the RBD-NP/AS03 immunization induced a geometric mean titer (GMT) of 1:63 on day 21 (3 weeks after primary immunization) that increased to 1:2,704 (43-fold) on day 42. The other groups, O/W, AS37, CpG-Alum, and Alum induced a GMT of 1:232, 1:640, 1:2,164, and 1:951 on day 42, respectively. These responses were remarkably higher than the nAb titers of 4 convalescent human samples (GMT 1:76) and the NIBSC control reagent (NIBSC code 20/130, nAb titer 1:241) (Extended Data Fig. 3a) assayed simultaneously. Next, we measured the nAb responses against the authentic SARS-CoV-2 virus using a recently established Focus Reduction Neutralization Titer (FRNT) assay^8^, which has been used to analyze the recent clinical trials of the Moderna mRNA vaccine^9,10^. Consistent with the pseudovirus neutralization assays, all adjuvants induced robust live-virus nAb titers after the secondary immunization (Fig. 1d). The RBD-NP/AS03 group showed the highest nAb titers (GMT 1:4,145) followed by the rest of the adjuvants with no statistical difference except for O/W. Furthermore, there was a strong correlation between pseudovirus and live-virus nAb titers, as seen in other studies (Extended Data Fig. 3b)^11,12^. Lastly, we measured the RBD-NP-specific plasmablast response using ELISPOT four days after secondary immunization (Extended Data Fig. 3c). The magnitude of antigen-specific IgG-secreting cells in blood correlated with the observed antibody responses (Extended Data Fig. 3d).

## RBD-NP/AS03 immunization induces durable live-virus nAb responses

Inducing potent and durable immunity is critical to the success of a vaccine and determines the frequency with which booster immunizations need to be administered. To determine durability of nAb responses, we followed five animals immunized with RBD-NP/AS03 without challenge for 5 months. The pseudovirus nAb titers measured until day 126 declined moderately but did not differ significantly between days 42 and 126 (Extended Data Fig. 4a). Strikingly, nAb response measured against the authentic SARS-CoV-2 virus using FRNT assay was durably maintained up to day 154 (Fig. 1e). Of note, the FRNT assay was performed in the same laboratory that measured durability in the Moderna vaccine study^10^. The GMT titers decreased by 5-fold between days 42 (GMT 5.638 in the 5 animals that were followed) and 154 (GMT 1,108), although this was not statistically significant (Fig. 1e). Furthermore, we observed little to no reduction in the efficiency of ACE-2 blocking by sera collected at these time points (Extended Fig. 4b). These results demonstrate that the RBD-NP/AS03 immunization induces potent and durable nAb responses^13^.

## Adjuvanted RBD-NP immunization elicits nAb response against the variant B.1.1.7

Variants of SARS-CoV-2 have been emerging recently, causing concerns that vaccine-induced immunity may suffer from lack of ability to neutralize the variants. One of the variants, B.1.1.7, was first identified in the United Kingdom and has since been found to be circulating globally. We evaluated if sera from animals immunized with RBD-NP + AS03, AS37, or CpG-Alum, neutralizes the B.1.1.7 variant. Using a pseudovirus neutralization assay as well as the live-virus neutralization assay, we determined that all the three groups induced nAb titers against the variant comparable to that of the wild-type (WT) SARS-CoV-2 (Extended Data Fig. 4c and Fig. 1f).

## Adjuvanted RBD-NP immunization induces robust CD4 T cell responses

We assessed antigen-specific T cell responses by intracellular cytokine staining (ICS) assay using a 21-parameter flow cytometry panel (Supplementary Table. 2). We first measured RBD-specific T cells after *ex vivo* stimulation with a peptide pool (15-mer peptides with 11-mer overlaps) spanning the SARS-CoV-2 RBD. RBD-NP immunization induced an antigen-specific CD4 T cell response but limited CD8 T cell response. RBD-specific CD4 responses were highest in the AS03 and CpG-Alum groups (Fig. 2a, b), and were significantly enhanced after secondary immunization. These responses were dominated by IL-2 or TNF-α-secreting CD4 T cells (Extended Data Fig. 5a), which remained detectable at day 42 (3 weeks post-secondary immunization). The median frequencies of IL-2^+^ and TNF-α^+^ CD4 T cell responses in the AS03 group were 0.1% and 0.08%, respectively, on day 28 and reduced to ∼0.07% on day 42. There was also a low but detectable IL-4 response in both the AS03 and CpG-Alum groups that peaked on day 28 but declined nearly to baseline levels by day 42 (Fig. 2b). Next to AS03 and CpG-Alum groups, Alum also induced a potent CD4 T cell response. Whereas 75% and 50% of animals in the Alum and O/W groups showed induction of RBD-specific CD4 T cells, respectively, the TLR-7 agonist AS37 induced a weak T cell response despite inducing potent antibody response in all the animals.

**Fig. 2.**
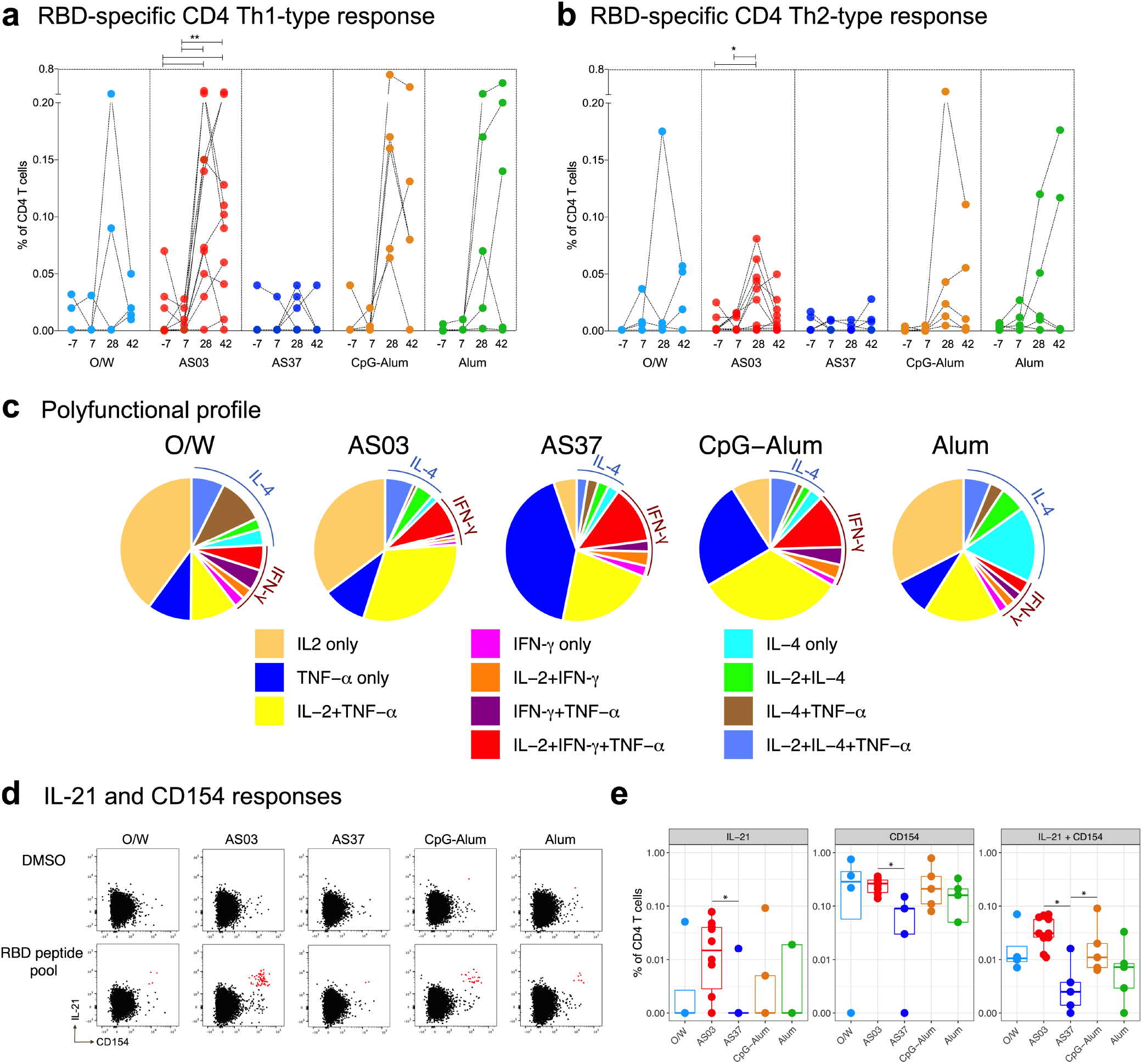
Cell-mediated immune responses to SARS-CoV-2 RBD-NP immunization. **a–b**, RBD-specific CD4 T cell responses measured in blood at time points indicated on the x axis. CD4 T cells secreting IL-2, IFN-γ, or TNF-α were plotted as Th1-type responses (**a**) and the Th2-type responses show the frequency of IL-4-producing CD4 T cells (**b**). **c**, Pie charts representing the proportions of RBD-specific CD4 T cells expressing one, two, or three cytokines as shown in the legend. **d**, Flow cytometry plots showing expression of IL-21 and CD154 after *ex vivo* stimulation with DMSO (no peptide, top) or an overlapping peptide pool spanning the SARS-CoV-2 RBD (bottom). **e**, RBD-specific CD154^+^ ± IL-21^+^ CD4^+^ T cell responses measured in blood at day 28. Asterisks represent statistically significant differences. The differences between groups were analyzed by two-sided Mann-Whitney rank-sum test and the differences between time points within a group were analyzed by two-sided Wilcoxon matched-pairs signed-rank test (* p < 0.05, ** p < 0.01).

We assessed the polyfunctional profile of antigen-specific CD4 T cells expressing IL-2, IFN-γ IL-4, and TNF-α (Fig. 2c). Although IL-2^+^, TNF-α^+^, and IL-2^+^TNF-α^+^ double-positive cells formed the majority (∼70%) in all adjuvant groups, differences between the groups were apparent. In particular, AS03 elicited similar proportions of polyfunctional Th1-type and Th2-type CD4 T cells, a balanced Th1/Th2 profile, CpG-Alum showed a slightly higher Th1-type response, and Alum a higher Th2-type response. We further extended our analyses to measure IL-21 and CD154, markers of circulating T_FH_-like cells for their critical role in germinal center formation and generation of durable B cell responses. We observed detectable IL-21 responses in the AS03 and CpG-Alum groups (Fig. 2d). All cells secreting IL-21 were CD154^+^. The IL-21^+^CD154^+^ double-positive cells were significantly higher in the AS03 and CpG-Alum groups in comparison with the AS37 group (Fig. 2e).

We also stimulated PBMCs with a peptide pool spanning the I53-50A and I53-50B nanoparticle component sequences to determine if RBD-NP immunization induces T cells targeting the nanoparticle scaffold. We observed a significant proportion of CD4 T cells targeting the I53-50 subunits with a response pattern, including polyfunctional profiles, similar to that of the RBD-specific T cells (Extended Data Fig. 5b,c). The frequencies of NP-specific CD4 T cells were ∼3-fold higher than that of RBD-specific CD4 T cells (Extended Data Fig. 5d), an observation that is consistent with the RBD making up approximately one third of the total peptidic mass of the immunogen. In summary, the RBD-NP immunization with adjuvants induced vaccine-specific CD4 T cells of varying magnitude. While IL-2 and TNF-α were the major cytokines induced by antigen-specific CD4 T cells, we also observed IL-21 and CD154 responses.

## RBD-NP immunization with different adjuvants protects NHPs from SARS-CoV-2 challenge

The primary endpoint of the study was protection against infection with SARS-CoV-2 virus, measured as a reduction in viral load in upper and lower respiratory tracts. To this end, we challenged the animals four weeks post-secondary immunization with 3.2 × 10^6^ PFU units via intratracheal and intranasal (IT/IN) routes. Viral replication was measured by subgenomic PCR quantitating the *E* gene RNA product on the day of the challenge, as well as 2-, 7- and 14-days post-challenge in nares, pharynges and BAL fluid.

Two days after challenge, 4 out of 4 control animals had detectable subgenomic viral RNA (*E* gene, range 3.1×10^5^ – 3.5×10^8^ viral copies) in the pharyngeal and the nasal compartments. By day 7, the viral RNA quantities reduced to baseline, consistent with previous studies^14,15^. All adjuvanted groups, except O/W, afforded protection from infection (Fig. 3a, b). In particular, none of the five animals challenged in the AS03 group had detectable viral RNA in pharyngeal swabs at any time and only one animal had detectable viral RNA in nasal swabs, at a level ∼1,000-fold lower than the median in control animals (2.2×10^4^ vs. 2.5×10^7^ viral copies). In contrast, viral RNA was detectable in pharyngeal swabs from all four animals in the SWE group, albeit at lower levels than the control group, and three out of four animals had detectable viral RNA in nasal swabs. Only one out of five animals in the CpG-Alum group had detectable viral RNA in pharyngeal or nasal swabs. The AS37 group and, remarkably, the Alum group also showed undetectable viral RNA in 3 of the 5 animals in both compartments (Fig. 3c).

**Fig. 3.**
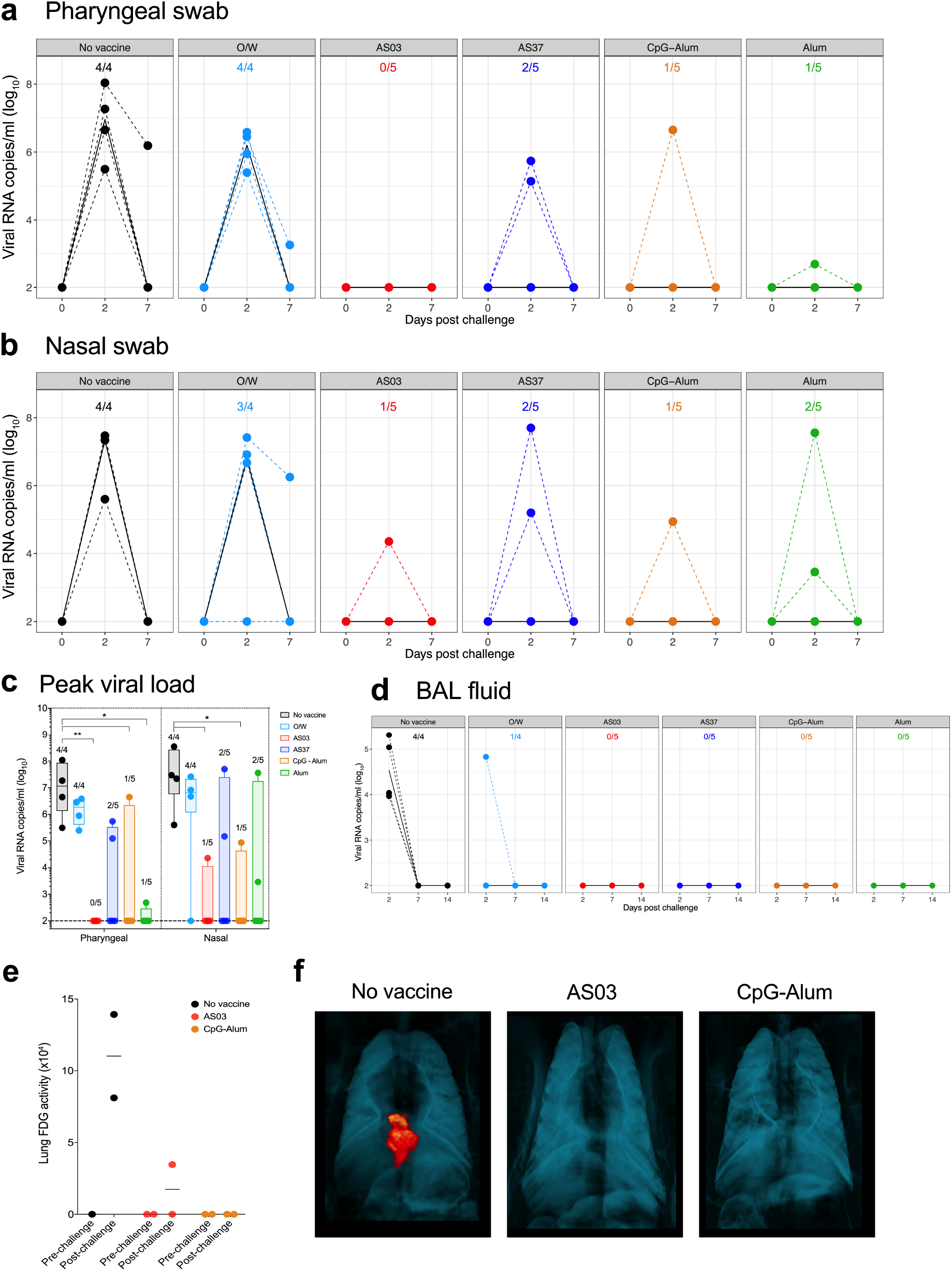
Protection against SARS-CoV-2 challenge. **a–b**, SARS-CoV-2 viral load in pharynges (**a**) and nares (**b**) of vaccinated and control macaques measured using subgenomic *E* gene PCR. **c**, Peak (day 2) viral load in pharyngeal and nasal compartments in each group. **d**, Viral load in BAL fluid measured using subgenomic *N* gene PCR. **e**, Inflammation in the lungs of two animals from each group indicated in the legend, pre-challenge (day 0) and post-challenge (day 4 or 5 after infection), measured using PET-CT scans. **f**, Representative PET-CT images of lungs from one animal in each group. In **a, b**, and **d**, the numbers within each box denote the number of infected animals per total number of animals in each group. PET signal is scaled 0 to 15 SUV. Statistical differences between groups were measured using two-sided Mann-Whitney rank-sum tests (* p < 0.05, ** p < 0.01).

We measured the subgenomic viral RNA in bronchoalveolar lavage (BAL) fluid to assess protection in the lung. We used a more sensitive PCR assay measuring the *N* gene product^16^ as we found only 2 control animals showing a positive viral load in the BAL using *E* subgenomic RNA. Two days after the challenge, all 4 of the four control animals showed a viral load in the range of 10^4^ - 10^6^ viral copies. In contrast, none of the animals in the vaccinated groups except one animal in the O/W group showed any detectable virus (Fig. 3d), suggesting effective protection in the LRT of all vaccinated groups, including the O/W group. Overall, the RBD-NP vaccination with adjuvants offered varying degree of protection against SARS-CoV-2 challenge in upper and lower respiratory tracts.

Vaccine-associated enhanced respiratory disease (VAERD) has previously been described for respiratory infections with respiratory syncytial virus and SARS-CoV^17,18^. Eosinophilia and enhanced inflammation in the lung have been shown to be associated with VAERD. We evaluated inflammation in the lung tissues of a subset of animals using PET-CT on the day of the challenge and 4 – 5 days post-challenge. Of the six animals evaluated (2 from no vaccine, 2 from AS03, and 2 from CpG-Alum groups selected randomly), we found inflammation in both control animals on day four compared to baseline, as measured by enhanced 2-Deoxy-2-[18F]fluoroglucose (FDG) uptake. In contrast, only one of the four vaccinated animals showed FDG uptake, to a much lesser extent than the control animals (Fig. 3e and Extended Data Fig. 6). These data are consistent with an absence of VAERD in these animals and suggest vaccination may prevent lung damage following SARS-CoV-2 challenge.

## Immune correlates of protection

Next, we set out to identify immune correlates of protection. Since we had five different adjuvant groups showing different protection levels within each group, we analyzed the correlations by combining animals from all the groups. We correlated humoral and cellular immune responses measured at peak time points (day 42 for antibody responses and day 28 for T cell responses) with the viral load (nasal or pharyngeal) to determine the putative correlates of protection in an unbiased approach. Neutralizing, both live and pseudovirus, titers emerged as the top statistically significant correlates of protection (Fig. 4a, b, and Extended Data Fig. 7a) in both nasal and pharyngeal compartments. Interestingly, NP-specific IL-2^+^ TNF^+^ CD4 T cell response also emerged as a statistically significant correlate of protection in both compartments (Fig. 4a and Extended Data Fig. 7b), the frequencies of which positively correlated with nAb titers (Extended Data Fig. 7c). This is consistent with the possibility that NP-specific CD4 T cells could offer T cell help to RBD-specific B cells.

**Fig. 4.**
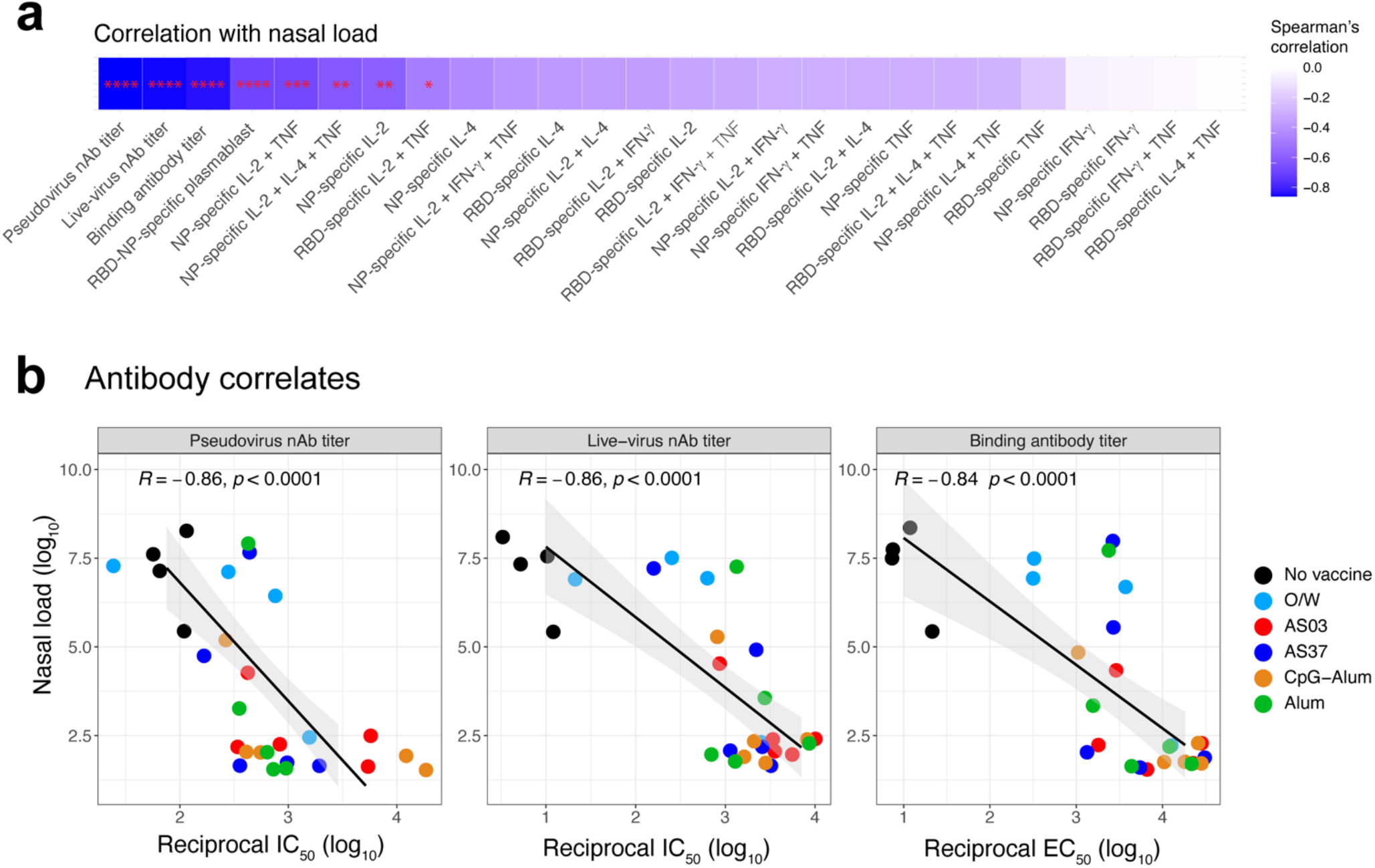
Immune correlates of protection. **a**, Heatmap showing Spearman’s correlation between peak nasal viral load (day 2) and various immune analyses readouts. All measurements were from peak time points (day 42 for antibodies, day 25 for plasmablast, and day 28 for T cell responses). The p-values were calculated for Spearman’s correlation and corrected for multiple-testing. Asterisks represent statistical significance. **b**, Spearman’s correlation plots between peak nasal viral load and the top three immune parameters shown in **a**.

## Systems serology profiling reveals functional antibody responses to RBD-NP vaccination

In addition to characterizing nAb and T cell responses to vaccination, we sought to understand the humoral functional profile elicited by each adjuvant. Vaccines rapidly induced a humoral immune response against SARS-CoV-2 spike with a profound increase in different anti-spike antibody isotypes (Fig. 5a - c) and FcR-binding (Fig. 5d, e) at day 21 and day 42. To understand how differences in the humoral response may lead to viral breakthrough, we performed a partial least square discriminant analysis (PLSDA) on the antibody features measured at day 42, using least absolute shrinkage and selection operator (LASSO) to select features to prevent overfitting (Fig. 5f). The PLSDA analysis showed separation between animals that had viral breakthrough in the nasal and pharyngeal and those that showed no viral breakthrough (Fig. 5f), marked by an enrichment in IgA, FcR3A and antibody-dependent neutrophil phagocytosis (ADNP) against spike in the protected animals (Fig. 5g).

**Fig. 5.**
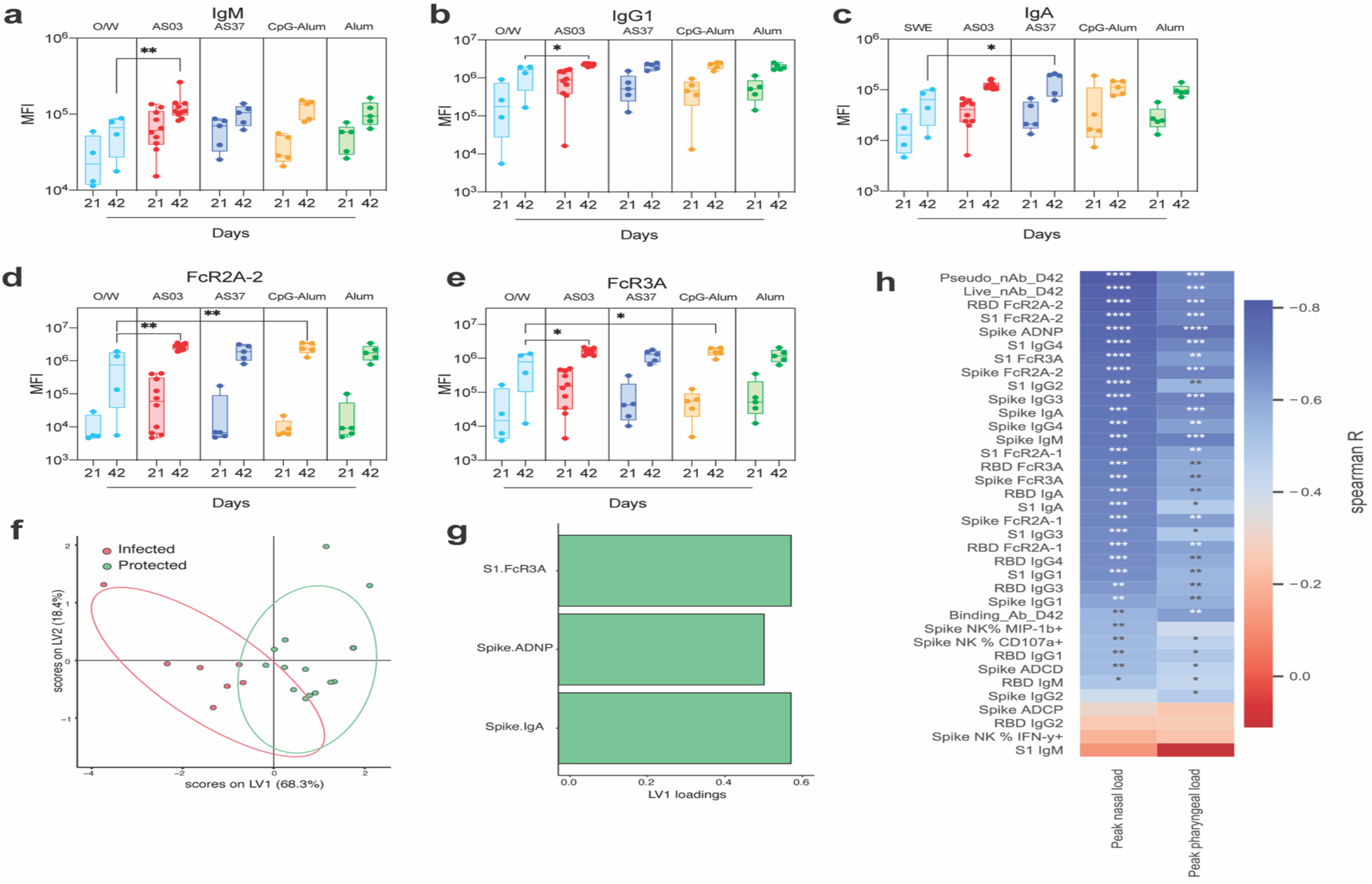
Functional antibody profiling by systems serology. **a–c**, SARS-CoV-2 Spike-specific binding IgM (**a**), IgG1 (**b**) and IgA (b) responses in sera collected at days 21 and 42. The box shows median and 25^th^ and 75^th^ percentiles and the error bars show the range. **d - e**, FcR-binding antibody responses, FcR2A-2 (**d**) and FcR3A (**e**) measured in serum collected at days 21 and 42. (**f**) PLSDA analysis of all antibody features measured using systems serology. (**g**) The top 3 antibody features discriminating protected vs. infected animals on day 42 in the PLSDA analysis. (**h**) Heatmap showing spearman’s correlation between peak nasal viral load (left) or pharyngeal vial load (right) and antibody responses (day 42) indicated on the Y-axis. The p-values were calculated for Spearman’s correlation and corrected for multiple-testing. In **a** - **e**, the statistically significant difference between two groups were determined by Mann-Whitney rank-sum test (* p < 0.05, ** p < 0.01, *** p < 0.001 and **** p < 0.0001).

We determined the correlation of each measured antibody feature and the peak nasal and pharyngeal viral load to further dissect the antibody features that provide protection against viral break-through. Whereas neutralizing Ab response still represents the strongest correlate of protection, we observed additional functional features including FcR binding (RBD FcR2A-2 and S1 FcR2A-2), and ADNP that were negatively correlated with nasal or pharyngeal viral loads (Fig. 5h) demonstrating a role for functional antibody responses in protection. Furthermore, each adjuvant group mounted a distinct profile of antibody response that correlated with protection against the virus (Extended Data Fig. 8). These differences between groups highlight that different adjuvants can elicit unique functional antibody responses to coordinate a protective antiviral response.

## Comparison of different Spike-based immunogens with AS03

The data described thus far demonstrate that RBD-NP immunogen when adjuvanted with AS03, AS37, CpG-Alum and Alum induce robust protective immunity. As a next step, we compared the immunogenicity of the RBD-NP immunogen to that of HexaPro, a highly stable variant of the prefusion Spike trimer^19^, in soluble or in a nanoparticle form. To this end, we designed a second study in which we immunized an additional 15 male RMs, distributed into three groups, with RBD-NP, soluble HexaPro or 20 Hexapro trimers displayed on the I53-50 nanoparticle (Hexapro-NP) (Fig. 6a, Supplementary table 3). All three groups were adjuvanted with AS03, the adjuvant that provided the highest magnitude of nAb responses and protection in all animals in the upper and lower respiratory tracts in previous experiments. The RBD-NP/AS03 immunization induced nAb titers comparable to that of the previous study, with a detectable titer on day 21 that boosted robustly at day 42. In comparison to the RBD-NP, soluble Hexapro or Hexapro-NP immunization induced notably higher nAb titers against the matched pseudovirus or authentic virus after one immunization (Fig. 6b, c). The RBD-NP, however, boosted strongly such that the magnitude of the nAb titers was not statistically different between the three groups on day 42 (Fig. 6b, c). Furthermore, the cross-reactive potential of the nAb response against the B.1.1.7 variant elicited by the soluble Hexapro immunization with AS03 was comparable to that of the WT virus (Fig. 6d), as was the case for RBD-NP (Fig. 1e). Taken together, these data indicate that the RBD-NP was as potent an immunogen as this highly stable version of the prefusion Spike trimer, consistent with previous observations that the vast majority of the neutralizing antibody response elicited by infection or immunization with trimeric Spike targets the RBD^13^. Moreover, these data suggest AS03 can be considered as a suitable adjuvant for clinical use with various forms of the Spike protein.

**Fig. 6.**
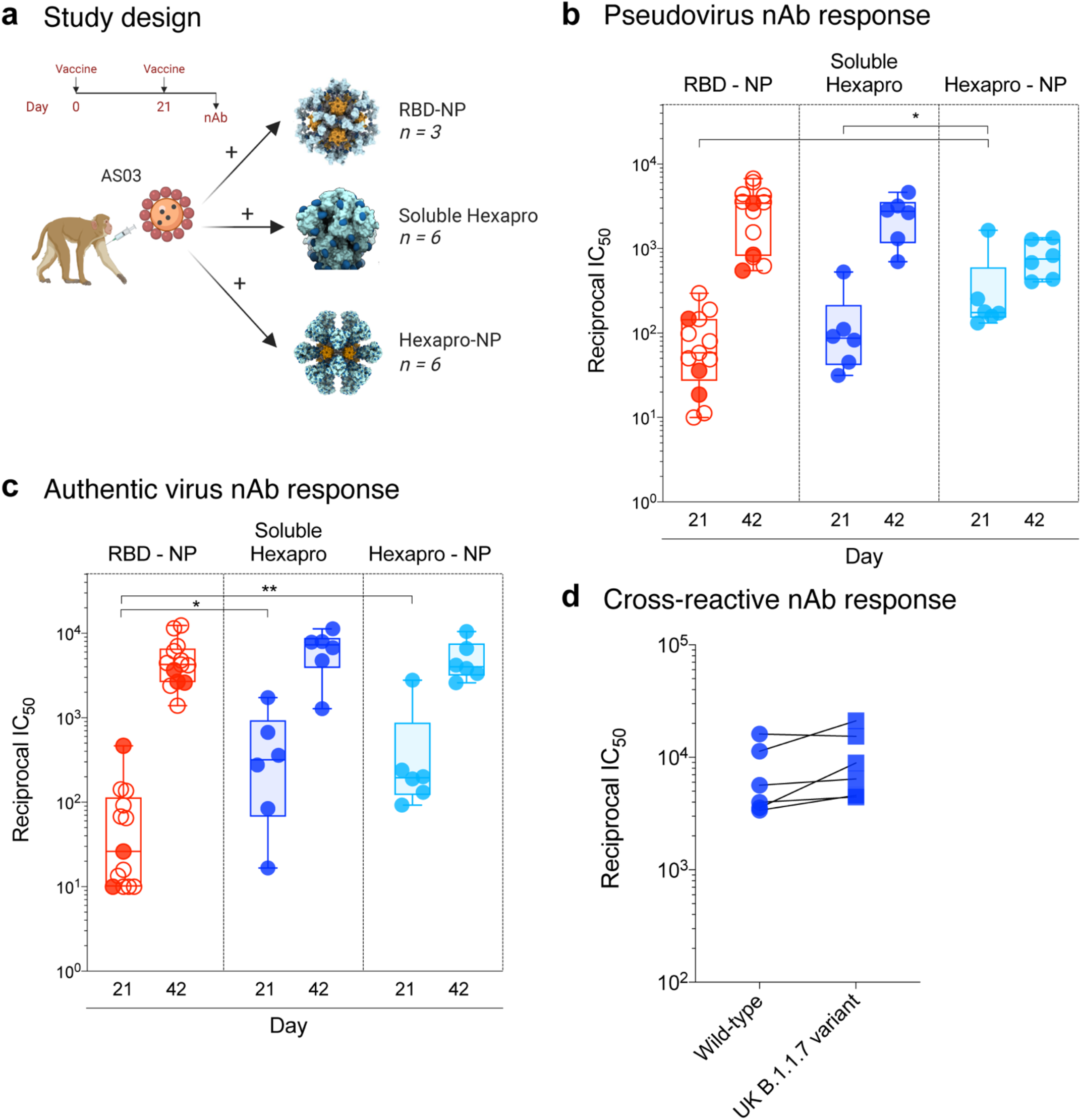
RBD-NP or HexaPro immunization with AS03 elicits comparable nAb responses. **a**, Schematic representation of the study design. **b – c**, Serum nAb titers (plotted as reciprocal IC_50_) determined using a SARS-CoV-2 S pseudovirus (**b**) or authentic SARS-CoV-2 (**c**) assay at day 21 and 42. The box shows median and 25^th^ and 75^th^ percentiles and the error bars show the range. Asterisks represent statistically significant differences between two groups analyzed by two-sided Mann-Whitney rank-sum test (* p < 0.05). Open circles denote animals from the earlier study shown in Fig. 1. **d**, Neutralizing antibody titers measured against live WT (circle) or B1.1.7. variant (squares) in sera collected on day 42 from animals that received soluble HexaPro.

## Discussion

The recent emergency use authorization of two messenger RNA (mRNA) vaccines against SARS-CoV-2 represents a major milestone in the fight against the COVID-19 pandemic^9,20-23^. However, manufacturing several billion doses of vaccines to vaccinate the entire world’s population will require a portfolio of different vaccine candidates. In particular, vaccinating special populations such as infants and the elderly could benefit from the use of subunit adjuvanted vaccine platforms with a demonstrable history of safety and efficacy in such populations^24,25^. The primary objective of this study was to select adjuvants for clinical development of the novel RBD-NP subunit vaccine candidate. We evaluated five different adjuvants, including two, Alum and AS03, that have been used in several millions of doses of licensed vaccines, for their capacity to elicit enhanced responses with the SARS-CoV-2 RBD-NP immunogen. All adjuvants tested induced substantial nAb titers (Fig. 1c, d). Of note, the nAb response to the authentic SARS-CoV-2 virus was induced to quantities equal to or higher than the titers observed in response to mRNA-1273 immunization in humans^9^ when measured using the same assay (FRNT assay) in the same laboratory. In general, all adjuvants induced detectable antigen-specific CD4 T cell responses, with AS03 and CpG-Alum inducing the highest frequencies. A notable finding was the induction of a high magnitude of CD4 T cell responses specific to the NP-scaffold. It is likely that these NP-scaffold-specific CD4 T cells could provide T cell help to RBD-specific B cells and promote B cell responses^26^.

Concomitant with these immune responses, we observed varying levels of protection against IN/IT challenge with SARS-CoV-2 virus in the various adjuvant groups. Furthermore, we did not observe any inflammation in the lungs 4 days post-challenge precluding the possibility of VAERD. The varying responses observed between the different groups allowed us to analyze putative correlates of protection. Using an unbiased correlation approach, we determined the nAb response as the primary correlate of protection; however, NP-specific IL-2^+^/TNF^+^ responses also showed a correlation. While this correlation is because of an indirect effect of correlation between T cells and nAb responses as showed in extended fig. 6C, or that the T cell represent an independent correlate of protection by synergizing with nAb response^27^ needs to be ascertained in future studies. Finally, and importantly, the nAb response induced by RBD-NP/AS03 immunization was durable. The nAb responses were measured by focus reduction neutralization test (FRNT) used in the longevity analysis of the Moderna vaccine candidate^10^. Although a direct comparison may not be ideal, the GMT IC_50_ in the FRNT assay on day 126 and 154 in this NHP study were 1,568 and 1,108, respectively, whereas the GMT IC_50_ on day 119 in response to mRNA-1273 was 775 in healthy young adults^10^. Finally, the adjuvanted RBD-NP immunization also induced cross-neutralization of the variant B.1.1.7.

In addition to evaluating clinically relevant adjuvants, we also compared the immunogenicity of RBD and prefusion-stabilized trimeric Spike immunogens. Our results demonstrate that the RBD-NP immunogen is as potent as immunogens based on the prefusion Spike trimer in inducing nAb titers. Whether differences in immunogenicity become apparent at lower doses of antigen warrants further investigation. Nonetheless, these data are encouraging as vaccine candidates in both antigenic formats (i.e., RBD vs. prefusion-stabilized trimeric Spike), each with distinct manufacturing considerations, move forward to the clinic. Of particular interest to the field will be to evaluate whether the nAb responses elicited by RBD-NP or HexaPro-based immunogens induces breadth not only against the new SARS-CoV-2 variants, but also against other coronaviruses.

Overall, the current study represents the most comprehensive comparative immunological assessment of a set of clinically relevant vaccine adjuvants and antigens in promoting robust and highly efficacious immune responses against a candidate subunit SARS-CoV-2 vaccine. These data reveal the promising performance of several adjuvants including AS03 and CpG 1018 (with Alum), which have been used in licensed vaccines, when used in conjunction with the SARS-CoV-2 RBD-NP immunogen. These results bode well for the clinical development of RBD-NP and other SARS-CoV-2 subunit vaccines with these adjuvants.

## Acknowledgements

This study was supported by the Bill & Melinda Gates Foundation INV – 018675 to B.P. and INV – 017592 to J.S.M.; OPP1156262 to D.V. and N.P.K., a generous gift from the Audacious Project, a generous gift from Jodi Green and Mike Halperin, a generous gift from the Hanauer family, the Defense Threat Reduction Agency (HDTRA1-18-1-0001 to N.P.K.), the National Institute of General Medical Sciences (R01GM120553 to D.V.), the National Institute of Allergy and Infectious Diseases (DP1AI158186 and HHSN272201700059C to D.V.; R01-AI127521 to J.S.M), a Pew Biomedical Scholars Award (D.V.), Investigators in the Pathogenesis of Infectious Disease Awards from the Burroughs Wellcome Fund (D.V.), Fast Grants (D.V.); and NIAID-NIH Contract: HHSN272201800004C to D.M. We thank the staff at both the New Iberia Research Center, UL Lafayette and the Tulane National Primate Research Center for the conduct of the animal studies. We also thank Dr. Lynda Stuart at the Bill and Melinda Gates Foundation for inputs and insights throughout the study.

## Author contributions

B.P., N.P.K., and H.K. conceptualized the study; B.P., P.S.A., F.V., N.P.K., and J.R. designed the study and were responsible for overall conduct of the study; S.W., D.P., M.C.M., E.K., C.S., N.B., M.M., B.F. and L.R. produced and purified RBD-NP and Hexapro-NP immunogens under the supervision of N.P.K.; J.F., and K.R organized and performed all macaque immunizations under the supervision of F.V.; L.S., D.E.F., and K.R. processed all samples collected during the immunization phase; P.S.A. and K.R. performed binding ELISA in Fig. 1 under the supervision of S.B.; A.C.W., and M.J.N, performed pseudovirus nAb response assays in Fig. 1 under the supervision of D.V.; V.V.E. and L.L. performed all authentic virus nAb assays under the supervision of M.S.; J,C.K. performed anti-Spike and anti-NP binding antibodies shown in Extended Fig. 2; P.S.A., C.L., and M.T. performed T cell assays; N.G., and P.A., organized all challenge experiments. N.G., P.A., K.R., J.D., L.D., R.B.B. and N.J.M. performed all post-challenge experiments; J.A.P., K.S.P., and C.R. provided challenge virus; A.W., and J.L.F. analyzed PET-CT data; P.S.A., and S.G. performed immune correlates analysis under the supervision of S.S.; C.A., S.F., A.Z., M.J.Z., and S.S. performed, analyzed and prepared figures of Systems serology under the supervision of G.A.; C-L.H. and J.S.M. provided Hexapro; X.S., and D.M., measured pseudovirus nAb response against UK B.1.1.7 variant; D.T.O., R.V.D.M., and R.R. provided AS03 and AS37 and guided formulation with the two adjuvants; R.L.C., and D.N., provided and guided CpG 1018 and its formulation with Alum; H.K. provided guidance throughout the project. P.S.A., and B.P. were responsible for the formal analysis of all datasets and preparation of figures; P.S.A., and B.P. wrote the manuscript with suggestions and assistance from all co-authors. All the authors read and accepted the final contents of the manuscript.

## Conflicts of interest

D.T.O., R.V.D.M., and R.R. are employees of GSK group of companies. R.L.C. and D.N. are employees of Dynavax Technologies Corporation. H.K. is an employee of Bill and Melinda Gates Foundation. C.-L.H. and J.S.M. are inventors on U.S. patent application no. 63/032,502 “Engineered Coronavirus Spike (S) Protein and Methods of Use Thereof.

**Extended Data Fig. 1.**
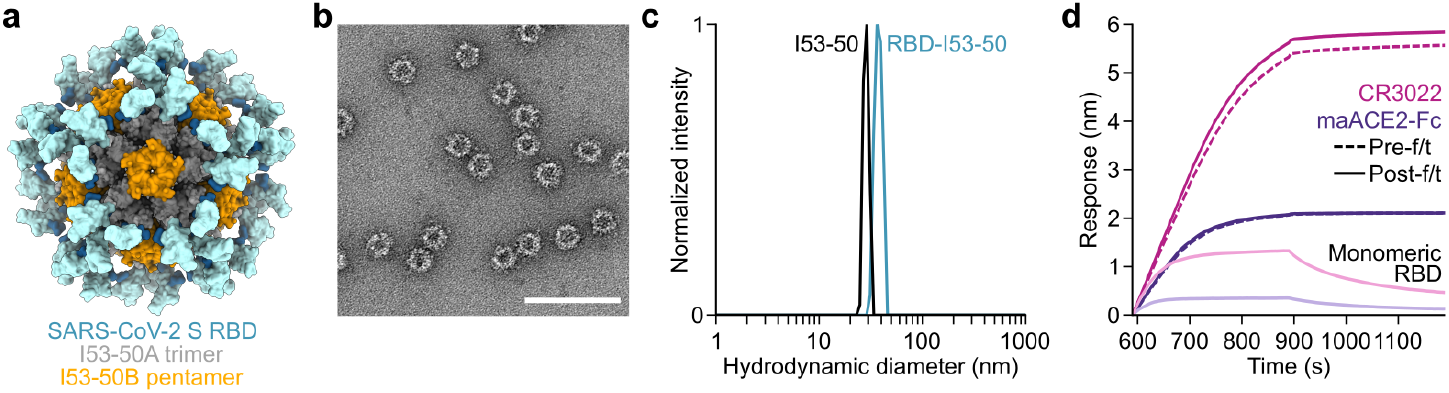
Structural, biophysical, and antigenic characterization of RBD-16GS-I53-50. **a**, Structural model of the RBD-16GS-I53-50 (RBD-NP) immunogen. The genetic linker connecting the RBD antigen to the I53-50A trimer is expected to be flexible and thus the RBD may adopt alternate orientations to that shown. **b**, Negative stain electron microscopy of RBD-NP. Scale bar, 100 nm. **c**, Dynamic light scattering (DLS) of RBD-NP and unmodified I53-50 lacking displayed antigen. The data indicate the presence of monodisperse nanoparticles with size distributions centered around 36 nm for RBD-NP and 30 nm for I53-50. In **b** and **c**, the samples were analyzed following a single freeze/thaw cycle. **d**, Antigenic characterization by biolayer interferometry (BLI). RBD-NP was bound to immobilized CR3022 mAb and maACE2-Fc receptor, both before and after one freeze/thaw cycle. Monomeric SARS-CoV-2 RBD was used as a reference antigen.

**Extended Data Fig. 2.**
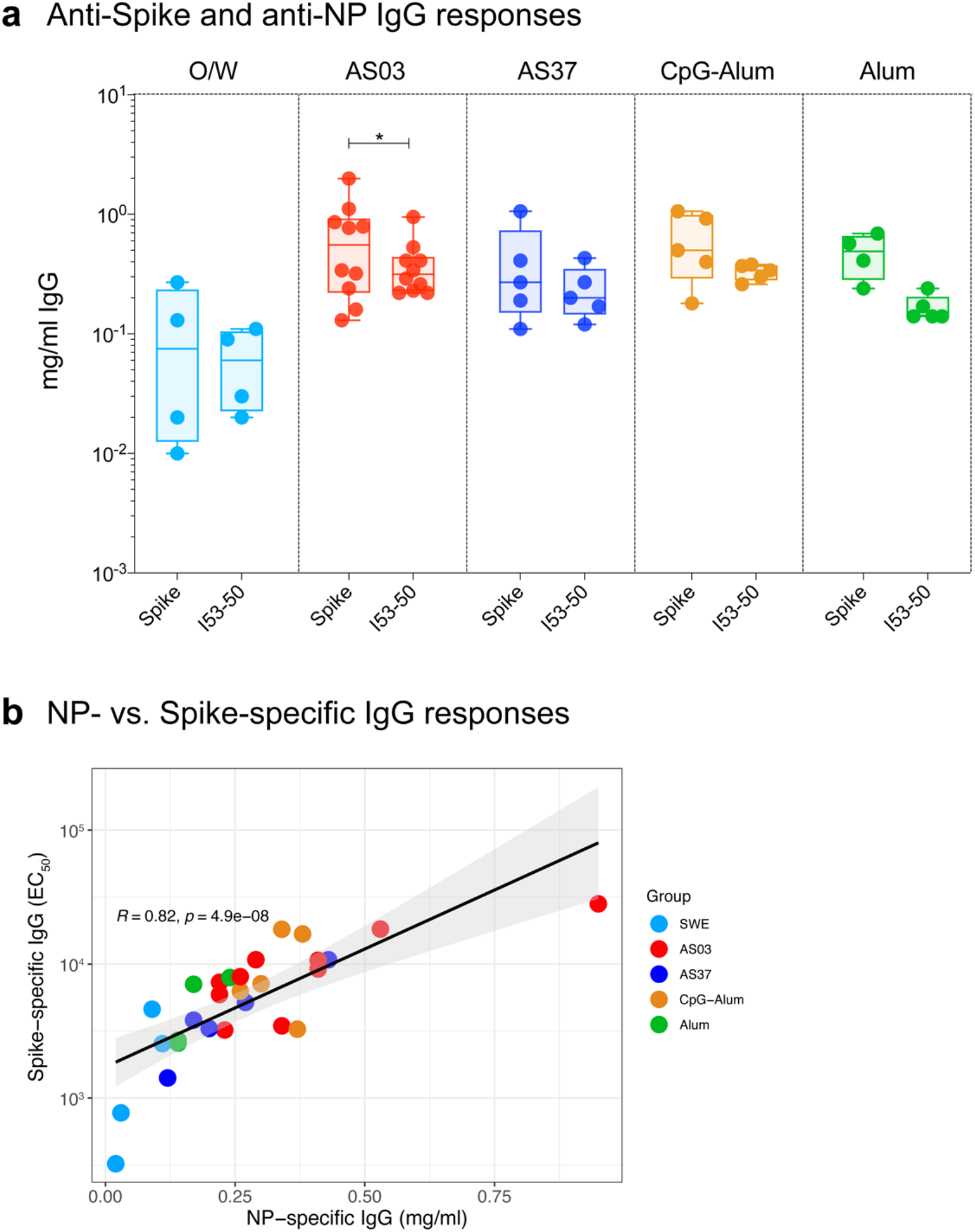
Comparison of anti-SARS-CoV-2 spike vs. anti-I53-50 nanoparticle scaffold antibody responses. **a**, Serum concentrations of anti-Spike IgG and anti-I53-50 nanoparticle IgG (anti-I53-50) in individual NHPs detected by ELISA at day 42. Boxes show median and 25^th^ and 75^th^ percentiles and the error bars show the range. The statistical difference between anti-Spike and anti-I53-50 IgG response was determined using two-sided Wilcoxon matched-pairs signed-rank test (* p < 0.05). **b**, Spearman’s correlation between anti-Spike IgG (described in Fig. 1) and anti-I53-50 IgG responses at day 42.

**Extended Data Fig. 3.**
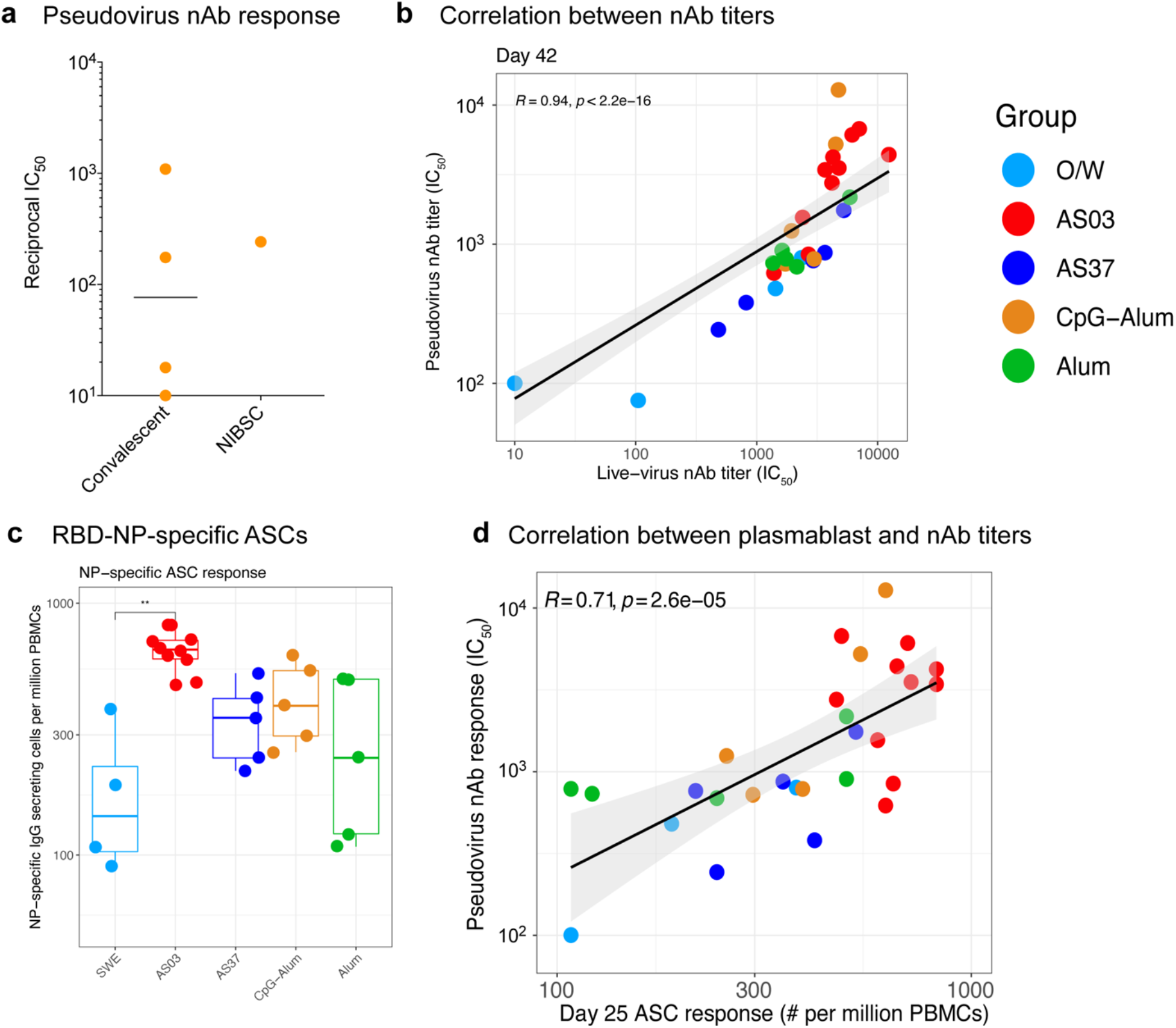
Humoral immune responses. **a**, Pseudovirus nAb response against human convalescent sera from 4 COVID-19 patients. **b**, Spearman’s correlation between pseudovirus and authentic virus nAb titers measured at day 42. **c**, RBD-NP-specific IgG secreting plasmablast response measured at day 4 post-secondary vaccination using ELISPOT. The difference between groups was analyzed using two-sided Mann-Whitney rank-sum test (** p < 0.01). **d**, Spearman’s correlation between plasmablast response on day 25 and pseudovirus nAb titer measured at day 42.

**Extended Data Fig. 4.**
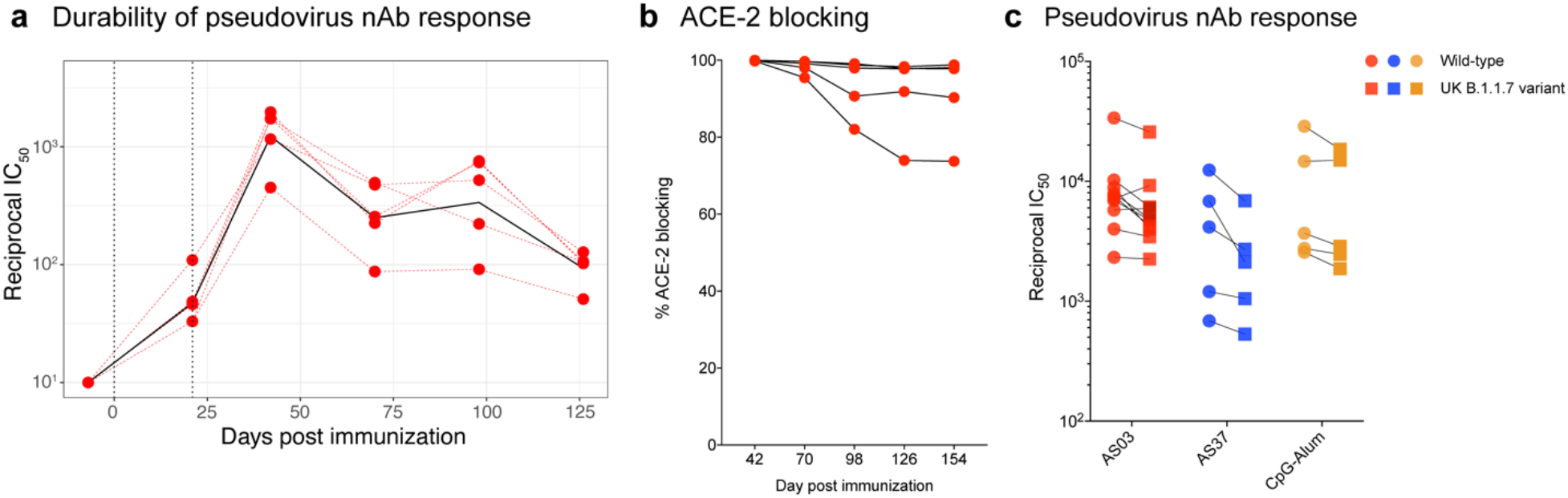
Durability and cross-neutralization. **a**, Pseudovirus nAb response measured in the AS03 durability group at time points indicated in X-axis. **b**, ACE-2 blocking measured in sera collected at time points indicated on the X-axis. **c**, SARS-CoV-2 nAb titers against pseudovirus wild-type containing D641G mutation on the Wuhan-1 Spike (circles) or the B.1.1.7 variant (squares) strain measured in day 42 sera.

**Extended Data Fig. 5.**
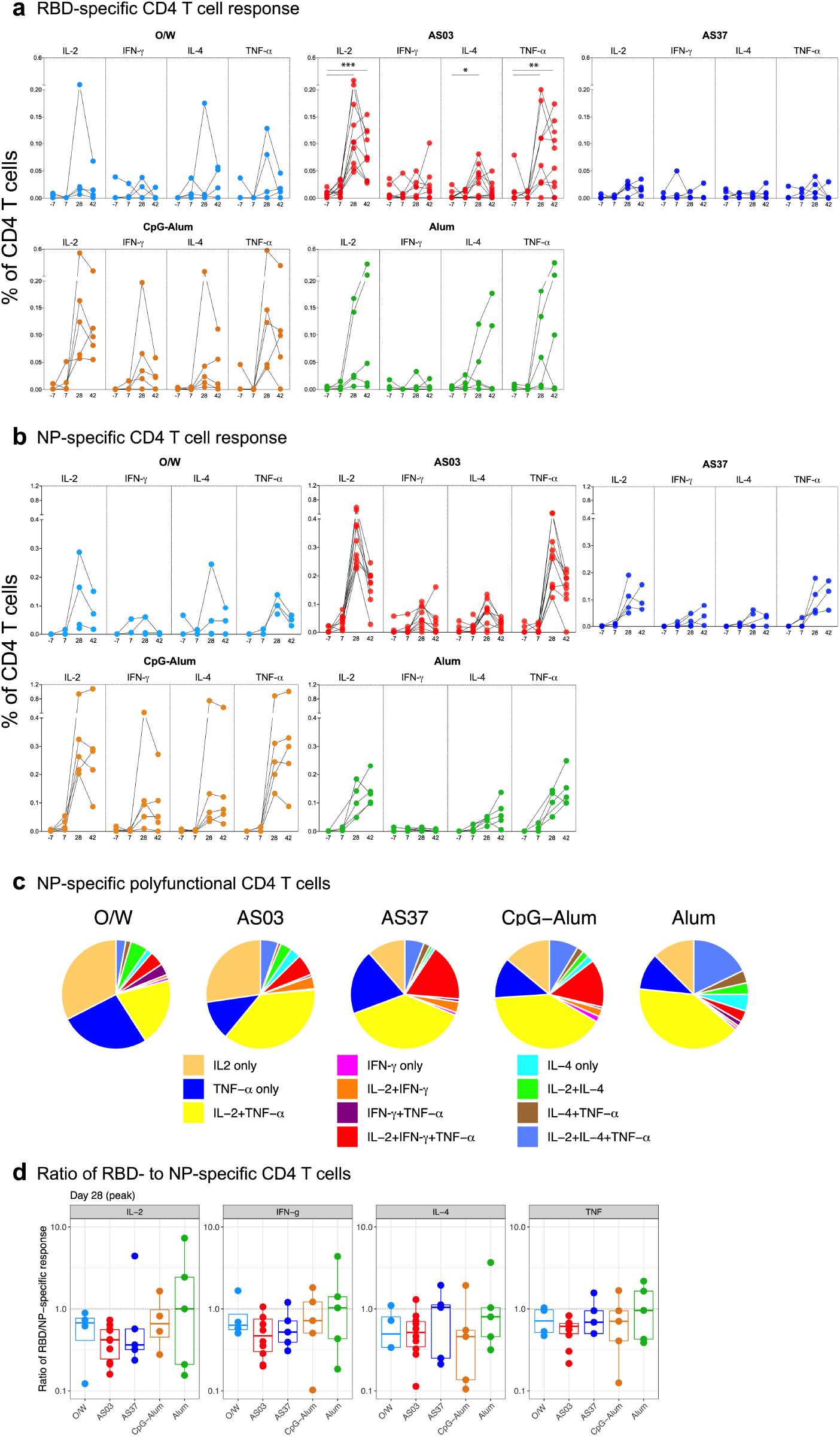
Cell-mediated immune responses to RBD-NP immunization. **a–b**, RBD- and NP-specific CD4 T cell responses measured in blood at time points indicated on the x axis. **c**, Pie charts representing the proportions of NP-specific CD4 T cells expressing one, two, or three cytokines as shown in the legend. **d**, Ratio of frequencies of RBD-specific to NP-specific CD4 T cells expressing cytokines indicated within each box. Asterisks represent statistically significant differences. The differences between time points within a group were analyzed by two-sided Wilcoxon matched-pairs signed-rank test (* p < 0.05, ** p < 0.01).

**Extended Data Fig. 6.**
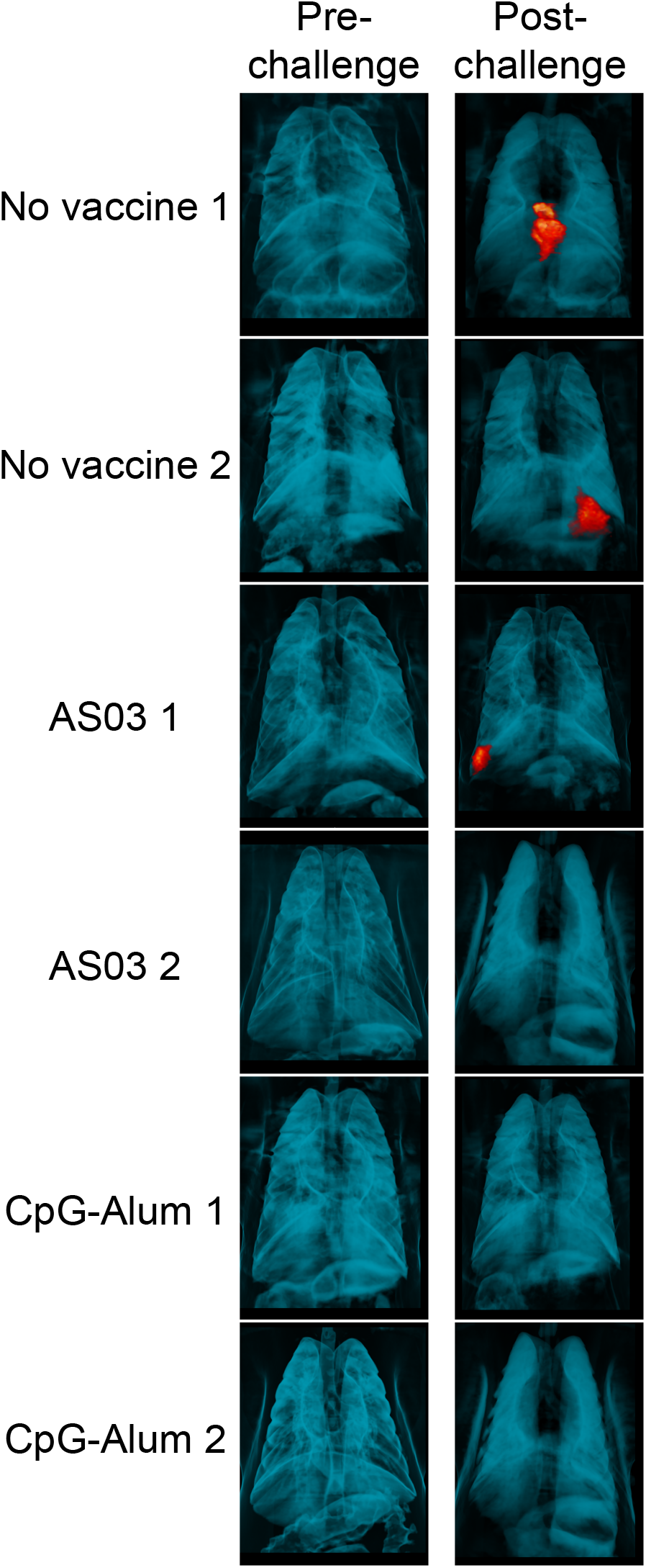
Inflammation in the lung. PET-CT images obtained from the lungs of SARS-CoV-2 infected animals from no vaccine, AS03, or CpG-Alum groups pre-challenge (day 0) and post-challenge (day 4 or 5).

**Extended Data Fig. 7.**
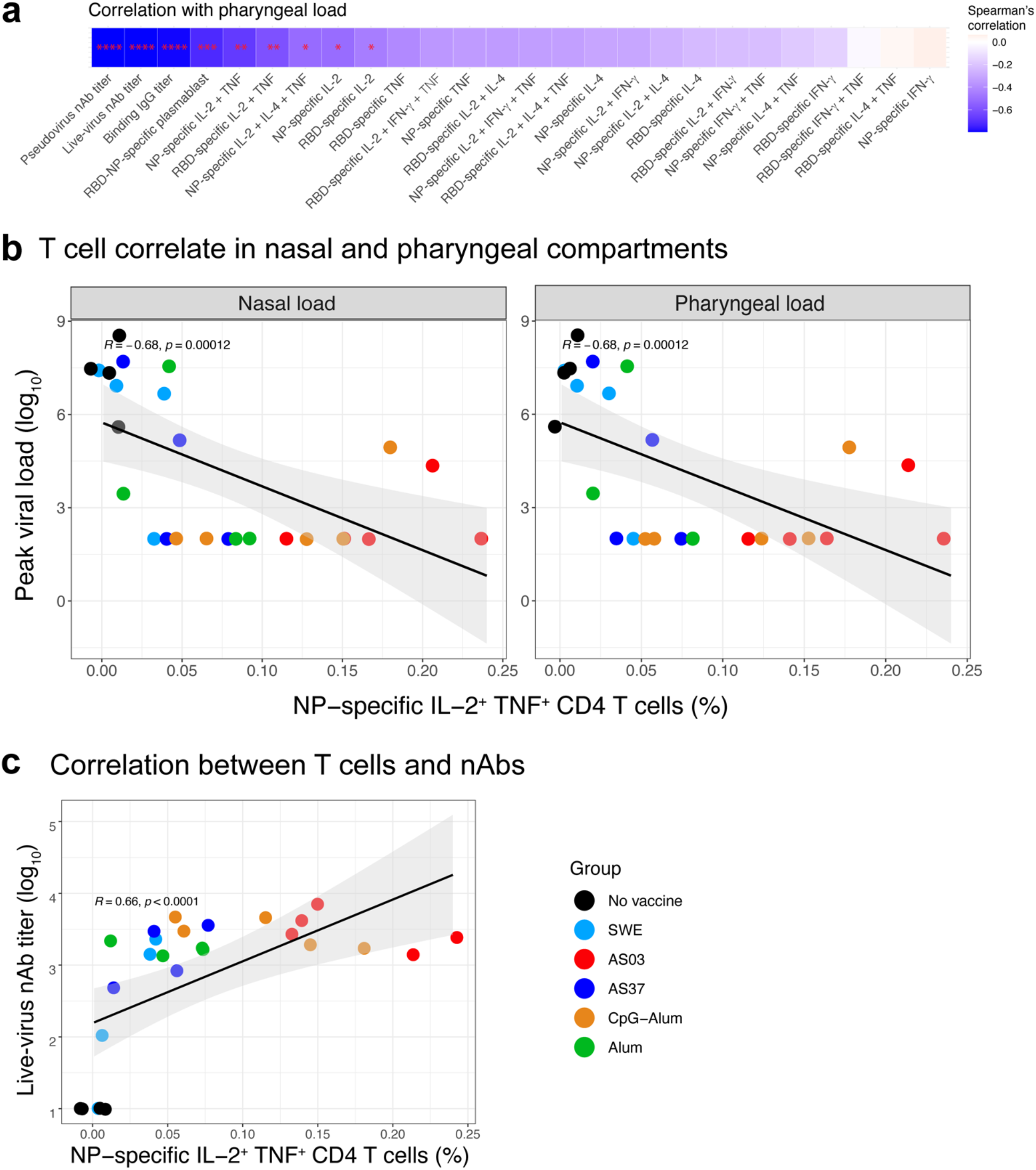
Immune correlates of protection. **a**, Heatmap showing Spearman’s correlation between peak pharyngeal viral load (day 2) and various immune parameters. All measurements were from peak time points (day 42 for antibodies, day 25 for plasmablast, and day 28 for T cell responses). The p-values were calculated for Spearman’s correlation and corrected for multiple-testing using the Benjamini-Hochberg method. **b**, Spearman’s correlation plots between peak nasal (left) or pharyngeal (right) viral load and the frequency of NP-specific IL-2^+^TNF-α^+^ CD4 T cells measured at day 28, 1 week after secondary immunization. **c**, Spearman’s correlation between the frequency of NP-specific IL-2^+^TNF-α^+^ CD4 T cells measured at day 28 and nAb response measured on day 42.

**Extended Data Fig. 8.**
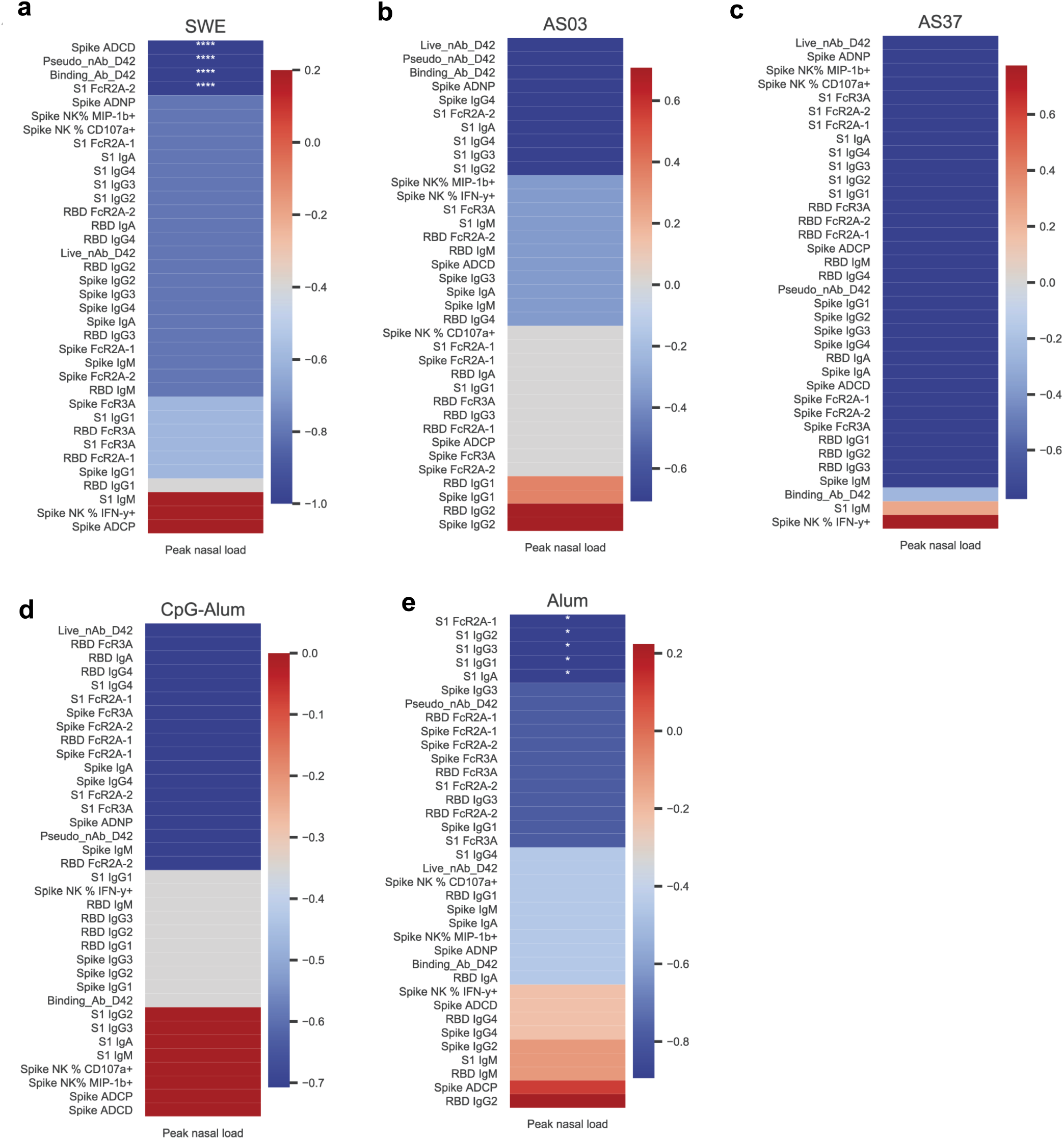
Antibody correlates of protection. Heatmap showing spearman’s correlation between peak nasal viral load (day 2) and antibody responses indicated on the Y-axis in groups of animals immunized with RBD-NP plus O/W (**a**), AS03 (**b**), AS37 (**c**), CpG-Alum (**d**) and Alum (**e**). The p-values were calculated for Spearman’s correlation and corrected for multiple-testing.

## Methods

### Animal subjects and experimentation

Thirty-three male rhesus macaques (*Macaca mulatta*) of Indian origin, aged 3 - 9 years were assigned to the study (Supplementary Table 1). Animals were distributed between the groups such that the age and weight distribution were comparable across the groups. Animals were housed and maintained as per National Institutes of Health (NIH) guidelines at the New Iberia Research Center (NIRC) of the University of Louisiana at Lafayette in accordance with the rules and regulations of the Committee on the Care and Use of Laboratory Animal Resources. The entire study (protocol 2020-8808-15) was reviewed and approved by the University of Louisiana at Lafayette Institutional Animal Care and Use Committee (IACUC). All animals were negative for SIV, simian T cell leukemia virus and simian retrovirus. For the challenge, the animals were transferred to the Regional Biosafety Level 3 facility at the Tulane National Primate Research Center, where the study was reviewed and approved by the Tulane University IACUC (Protocol 3918).

### RBD-16GS-I53-50 nanoparticle immunogen production

Nanoparticle immunogen components and nanoparticles were produced in the same manner as previously described in detail^5^, with the exception that the nanoparticle was in a buffer containing 50 mM Tris pH 8, 150 mM NaCl, 100 mM L-Arginine, 5% sucrose.

### Nanoparticle biochemical characterization

Dynamic light scattering, negative stain electron microscopy, and maACE2-Fc and CR3022 IgG biolayer interferometry were performed as described previously^5^.

### Adjuvant formulations and immunization

Essai O/W 1849101, a squalene-in-water emulsion (O/W) was kindly provided by Seppic. The Vaccine Formulation Institute (VFI) established the formulation of RBD-NP with O/W. For each dose, RBD-NP was diluted to 50 μg/ml (RBD component) in 250 μl of Phosphate buffered saline with 150 mM NaCl and mixed with an equal volume of O/W. The dose of O/W was 50% v/v. AS03 and AS37 were kindly provided by GSK Vaccines. AS03 is an oil-in-water emulsion that contains 11.86 mg α-tocopherol, 10.69 mg squalene, and 4.86 mg polysorbate 80 (Tween-80) in PBS, whereas AS37 is a TLR-7 agonist (200 μg/ml) adsorbed to Aluminium hydroxide (2 mg/ml). For each dose, RBD-NP was diluted to 50 μg/ml (RBD component) in 250 μl of Tris-buffered saline (TBS) and mixed with an equal volume of AS03 or AS37. The dose of AS03 was 50% v/v (equivalent of one human dose), AS37 included 50 μg TLR-7 agonist and 0.5 mg Aluminium hydroxide. CpG 1018 was generously provided by Dynavax Technologies at a concentration of 12 mg/ml. Alum (Alhydrogel 2%) was purchased from Croda Healthcare (Batch #0001610348). Of note, we used CpG-Alum in contrast to CpG 1018 (no Alum) used in Heplisav-B. For each dose of CpG-Alum, 25 μg antigen (RBD component) in TBS was mixed with 0.75 mg Alum and incubated on ice for 30 min. After 30 min of incubation, 1.5 mg of CpG 1018 was added and mixed rapidly. Each dose contained 1.5 mg CpG 1018 and 0.75 mg Alum. For each dose of Alum, 25 μg antigen (RBD component) in TBS was mixed with 0.75 mg Alum, matching the concentration of Alum in the CpG-Alum formulation, and incubated on ice for 30 min. Soluble Hexapro or Hexapro-NP used in study 1B was diluted to 50 μg/ml in 250 μl of Tris-buffered saline (TBS) and mixed with an equal volume of AS03. All immunizations were administered via the intramuscular route in right forelimbs. The volume of each dose was 0.5 ml.

### Anti-S binding ELISA

SARS-CoV-2 Spike protein was produced in HEK293T cells (Atum, Newark, CA). 96-well Corning Costar high binding plates (Thermo Fisher Scientific) were coated with SARS-CoV-2 Sike protein in phosphate-buffered saline (PBS) at a concentration of 0.2 µg per well overnight at 4°C. On the next day, wells were washed 3x with PBS - 0.1% Tween 20 (PBS-T) and blocked with PBS-T containing 3% non-fat milk powder for 1 hour at room temperature (RT). Wells were then incubated with plasma samples from NHPs at different dilutions starting at 1:100 in PBS-T containing 1% non-fat milk for 1 hour at 37°C. After washing 3x with PBS-T, horseradish peroxidase conjugated goat anti-monkey IgG (γ-chain specific, Alpha Diagnostics, 1:4’000 dilution), or IgA (α-chain specific, Alpha Diagnostics, 1:4’000 dilution) in PBS-T containing 1% non-fat milk was added and incubated for 1 hour at RT. Wells were washed 3x with PBS-T before addition of 3,3’,5,5’-Tetramethylbenzidine (TMB) substrate solution. The reaction was stopped after 12 minutes by addition of 0.16 M sulfuric or 1 M hydrochloric acid. The optical density (OD) at 450 nanometers was measured with a Tecan Infinite M Nano Plus microplate reader.

### Anti-I53-50 ELISA

The protocol was adapted from Tiller, et al.2008^28^. Briefly, recombinant I53-50 protein nanoparticles, SARS-CoV-2 S2P trimers, or goat anti-human IgG (Jackson ImmunoResearch, #109-005-044) were immobilized on 96-well Nunc MaxiSorp (Thermo Fisher Scientific) plates (2 μg/mL, 50 μL/well). After 1 h incubation at room temperature, plates were blocked with 200 μL TBS plus 2% (w/v) BSA and 0.05% (v/v) Tween20 for 1 h. Plates were washed 3x in TBST by plate washer (BioTek), and 50 μL of 1:5 serial dilutions starting at 1:100 of NHP sera in TBST incubated for 1 h in wells with I53-50 or spike. In wells with anti-human IgG capture antibody, human IgG control (SinoBiological, #HG1K) was serially diluted from 0.5-500 ng/mL in TBST in triplicate and 50 μL of each dilution incubated for 1 h. Plates were washed 3x in TBST, then HRP-conjugated goat anti-monkey IgG (Alpha Diagnostics, 70021) was diluted 1:5,000 in 2% BSA in TBST and 50 μL incubated in each well for 30 min. Plates were washed 3x in TBST and 100 μL of TMB (SeraCare) was added to each well for 2 min. The reaction was quenched by adding 100 μL of 1 N HCl. Plates were immediately read at 450 nm on a SpectraMax M5 plate reader (Molecular Devices) and data plotted and fit in Prism (GraphPad) using nonlinear regression sigmoidal, 4PL, X is log(concentration) to determine EC_50_ values from curve fits. A logarithmic equation fit to the linear portion of the sigmoidal curve of the human IgG control was used to calculate mg/mL of IgG in sera for anti-I53-50 and anti-Spike titers. All steps were performed at ambient temperature.

### Pseudovirus production and neutralization assay

Pseudovirus production has been described in Walls et al.2020^7^. Briefly, MLV-based SARS-CoV-2 Spike pseudotyped viruses were prepared as previously^7,29,30^ except that the SARS-CoV-2 S construct contained the D614G mutation and a truncation of the C-terminal 21 residues^12,31^.

For neutralization assays, HEK-hACE2 cells were cultured in DMEM with 10% FBS (Hyclone) and 1% PenStrep with 8% CO_2_ in a 37°C incubator on poly-lysine (sigma) 96 well plates. To coat plates, 40 μL of poly-lysine (Sigma) was incubated with rotation for 5 min. Poly-lysine was removed, plates were dried for 5 min then washed once with water prior to plating cells. The following day, cells were checked to be at 80% confluence. In a half-area 96-well plate a 1:3 serial dilution of sera was made in DMEM in 22 μL final volume. 22 μL of pseudovirus was then added to the serial dilution and incubated at room temperature for 30 - 60 min at room temperature. HEK-hACE2 plate media was removed and 40 μL of the sera/virus mixture was added to the cells and incubated for 2 h at 37°C with 8% CO_2_. Following incubation, 40 μL 20% FBS and 2% PenStrep containing DMEM was added to the cells. Following 48 - 72h infection, One-Glo-EX (Promega) was added to the cells in half culturing volume (40 μL added) and incubated in the dark for 5 min prior to reading on a Varioskan LUX plate reader (Thermo Fisher Scientific). Measurements were done on all sera samples from each group in at least duplicates. Relative luciferase units were plotted and normalized in Prism (GraphPad) using a zero value of cells alone and a 100% value of 1:2 virus alone. Nonlinear regression of log(inhibitor) vs. normalized response was used to determine IC_50_ values from curve fits.

### Focus Reduction Neutralization Titer assay

Neutralization assays with authentic SARS-CoV-2 virus were performed as previously described^32^. Plasma/serum were serially diluted (three-fold) in serum-free Dulbecco’s modified Eagle’s medium (DMEM) in duplicate wells and incubated with 100–200 FFU infectious clone derived SARS-CoV-2-mNG virus^33^ at 37 °C for 1 h. The antibody-virus mixture was added to VeroE6 cell (C1008, ATCC, #CRL-1586) monolayers seeded in 96-well blackout plates and incubated at 37 °C for 1 h. Post-incubation, the inoculum was removed and replaced with pre-warmed complete DMEM containing 0.85% methylcellulose. Plates were incubated at 37 °C for 24 h. After 24 h, methylcellulose overlay was removed, cells were washed twice with PBS and fixed with 2% paraformaldehyde in PBS for 30 min at room temperature. Following fixation, plates were washed twice with PBS and foci were visualized on a fluorescence ELISPOT reader (CTL ImmunoSpot S6 Universal Analyzer) and enumerated using Viridot^34^. The neutralization titers were calculated as follows: 1 - (ratio of the mean number of foci in the presence of sera and foci at the highest dilution of respective sera sample). Each specimen was tested in two independent assays performed at different times. The FRNT-mNG_50_ titers were interpolated using a 4-parameter nonlinear regression in GraphPad Prism 8.4.3. Samples with an FRNT-mNG_50_ value that was below the limit of detection were plotted at 10. For these samples, this value was used in fold reduction calculations.

### ACE-2 blocking assay

Antibodies blocking the binding of SARS-CoV-2 Spike RBD to the angiotensin-converting enzyme 2 (ACE2) were detected with a V-PLEX SARS-CoV-2 Panel 2 (ACE2) Kit (Meso Scale Diagnostics) according to the manufacturer’s instructions. Serum samples from NHPs were analyzed in duplicate at a dilution of 1:100 and percent inhibition was calculated based on the equation ((1 – Average Sample ECL Signal / Average ECL signal of Calibrator 7) x 100).

### Pseudovirus neutralization assay against UK B.1.1.7

Neutralization assay evaluating the ability of sera from vaccinated animals to neutralize wildtype (with D614G in spike) versus the B.1.1.7 variant viruses were performed using a pseudotyped virus neutralization assay previously reported with minor modifications^35^. Briefly, mutations were introduced into a plasmid expressing codon-optimized Spike of the Wuhan-1 strain that contains the D614G mutation using site-directed mutagenesis. Pseudovirions were produced in HEK293T/17 cells by co-transfection of a lentivirus backbone plasmid, a Spike-expressing plasmid, and a firefly Luc reporter gene plasmid. Pseudotyped viruses were titrated in 293T/ACE2.MF cells for TCID50 and used for neutralization assay. Virus were incubated with serial diluted serum samples at 37°C for 1 hr, and subsequent added to cells and incubated for 66-72 hrs. Luminescence was measured using a GloMax Navigator luminometer (Promega). Neutralization titers are the inhibitory dilution (ID) of serum samples at which RLUs were reduced by either 50% (ID50) or 80% (ID80) compared to virus control wells after subtraction of background RLUs.

### Intracellular cytokine staining assay

Antigen-specific T cell responses were measured using the ICS assay. Live frozen PBMCs were revived, counted and resuspended at a density of 1 million live cells/ml in complete RPMI (RPMI supplemented with 10% FBS and antibiotics). The cells were rested overnight at 37°C in CO_2_ incubator. Next morning, the cells were counted again, resuspended at a density of 15 million/ml in complete RPMI and 100 µl of cell suspension containing 1.5 million cells was added to each well of a 96-well round-bottomed tissue culture plate. Each sample was treated with three conditions, no stimulation, a peptide pool spanning the RBD region of spike at a concentration of 1.2 µg/ml of each peptide and a peptide pool spanning the I53-50A, and I53-50B components of the NP-scaffold (1.2 µg/ml of each peptide) in the presence of 1 µg/ml of anti-CD28 (clone CD28.2, BD Biosciences) and anti-CD49d (clone 9F10, BD Biosciences) as well as anti-CXCR3 and anti-CXCR5 (clone and concentration details in supplementary table 2). The peptides were custom synthesized to 90% purity using GenScript, a commercial vendor. All samples contained 0.5% v/v DMSO in total volume of 200 µl per well. The samples were incubated at 37°C in CO2 incubators for 2 h before addition of 10 µg/ml Brefeldin-A. The cells were incubated for an additional 4 h. The cells were washed with PBS and stained with Zombie UV fixable viability dye (Biolegend). The cells were washed with PBS containing 5% FCS, before the addition of surface antibody cocktail (Supplementary table 1). The cells were stained for 20 min at 4°C in 100 µl volume. Subsequently, the cells were washed, fixed and permeabilized with cytofix/cytoperm buffer (BD Biosciences) for 20 minutes. The permeabilized cells were stained with ICS antibodies for 20 min at room temperature in 1X-perm/wash buffer (BD Biosciences). Cells were then washed twice with perm/wash buffer and once with staining buffer before analysis using BD Symphony Flow Cytometer. All flow cytometry data were analyzed using Flowjo software v10 (TreeStar Inc.).

### Viral challenge

Animals were inoculated via the intratracheal (IT) and intranasal (IN) routes with a total of 3.2 × 10^6^ PFU of SARS-CoV-2, isolate USA WA1/2020 (Accession: MN985325). The virus stock was generated by expansion of a seed stock on Vero E6 cells and titered by plaque assay on Vero E6 cells. It was deep sequenced and found to contain no polymorphisms at greater than 5% of reads relative to the original patient isolate. The furin cleavage site, a site with frequent culture adaptation in Vero E6 cells, harbored no polymorphisms at greater than 1% of sequence reads in this stock.

### Sampling of nares and pharynges

The animals were anesthetized and placed in dorsal recumbency or a chair designed to maintain its upright posture. The pharynx was visualized using a laryngoscope. A sterile swab was gently rubbed/rolled across the lateral surfaces of the pharynx for approximately five seconds. The tonsillar fossa and posterior pharynx were included. Care was taken to avoid touching the soft palate, uvula, buccal mucosa, tongue, or lips. After all pertinent surfaces have been sampled, the swab is removed and placed into either culture medium or an appropriate container for transport. The pharyngeal swabs were done prior to the nasal swabs to reduce blood contamination from the nasal cavity down into the pharyngeal area.

Sterile swabs were gently inserted into the nares. Once inserted, the sponge/swab was rotated several times within the cavity/region and immediately withdrawn.

### Bronchoalveolar lavage (BAL) collection and processing

The animals were anesthetized using Telazol and placed in a chair designed specifically for the proper positioning for BAL procedures. A local anesthetic (2% lidocaine) may be applied to the larynx at the discretion of the veterinarian. A laryngoscope is used to visualize the epiglottis and larynx. A feeding tube is carefully introduced into the trachea after which the stylet is removed. The tube is advanced further into the trachea until slight resistance is encountered. The tube is slightly retracted and the syringe is attached. Aliquots of warmed normal saline are instilled into the bronchus. The saline is aspirated between each lavage before a new aliquot is instilled. When the procedure is complete, the animal is placed in right lateral recumbency. The animal is carefully monitored with observation of the heart rate, respiratory rate and effort, and mucous membrane color. An oxygen facemask may be used following the procedure at the discretion of the veterinarian. The animal is returned to its cage, positioned on the cage floor in right lateral recumbency and is monitored closely until recovery is complete.

The BAL samples were filtered twice via 100 μ strainers and collected in 50 ml centrifuge tubes. The samples were centrifuged at 300 g for 10 min at 4°C. The supernatant was transferred into new tubes, aliquoted and stored at −80°C until RNA isolation. The cells were washed, lysed for red-blood cells using ACK-lysis buffer and live-frozen in 90% FBS + 10% DMSO.

### Viral load

Quantitative RT-qPCR (reverse transcriptase - quantitative PCR) was performed as we described previously^36^. RT-qPCR for the subgenomic (sg) RNA encoding the Envelope (E) protein was performed as described^37^ and for the sgRNA encoding the Nucleocapsid (N) protein was performed using the same cycling conditions as used for the sg-E-RT-qPCR using an unpublished assay kindly provided by Drs. Dennis Hartigan-O’Connor and Joseph Dutra (U. California-Davis). Primers and probes for the sg N qRT-PCR were as follows; Forward 5’-CGATCTCTTGTAGATCTGTTCTC-3’, Reverse 5’-GGTGAACCAAGACGCAGTAT-3’, probe 5’-FAM-TAACCAGAATGGAGAACGCAGTGGG-BHQ1-3’. Both PCRs were run in volume of 20 μl containing 5 μl sample, 900 nM primers, 250 nM probe with TaqPath 1-step RT-qPCR master mix, CG (Thermo Fisher Scientific). The PCR conditions were 2 min at 25°C for UNG incubation, 15 min at 50°C for reverse transcription, 2 min at 95°C for Taq activation, followed by 40 cycles of 95°C, 3 s for denaturation and 60°C, 30 s for annealing and elongation.

### PET-CT administration, acquisition and data collection

The animals were anesthetized and brought to the PET-CT suite where they were monitored and prepared for imaging. An intravenous (IV) catheter is placed and the animals were intubated and placed on a gas anesthetic (isoflurane). 2-Deoxy-2-[18F]fluoroglucose (FDG) was administered as an IV bolus at a dose of 0.5 mCi/kg in the animal preparatory room. The catheter was flushed, and the animals were transferred to the PET/CT imaging room. Images were acquired on a Mediso LFER 150 PET/CT (Mediso Medical Imaging Systems, Budapest, Hungary). The animals were then placed on the table in a ‘head-in-supine’ position with heat support. Scout CT images of side and top views were obtained for positioning purposes and preferred scanning ranges. The number of fields of view (FOV) was determined depending on the size of the animal (each FOV covers 15 cm and takes 10 minutes to obtain with PET). A CT scan was captured at 80 kVp and 1 mA with a time range of 1-5 minutes depending on the FOV. Breath holds were performed during the CT scan on animals that can be imaged in one FOV. A breath-hold lasts for the majority of the CT scan which is approximately 45-60 seconds. PET images were obtained following FDG uptake time (45-60 minutes) and the CT scan. Once the images were captured, the animal’s fluids were discontinued and the animal was removed from isoflurane. When swallowing reflexes return, the animal was extubated and returned to its home cage. Images were reconstructed using Nucline software with the following parameters: Mediso Tera-Tomo 3D algorithm, 8 iterations, 9 subsets, voxel size 0.7 mm.

### PET-CT data analysis

PET-CT images were analyzed using OsiriX MD or 64-bit (v.11, Pixmeo, Geneva, Switzerland). Before analysis, the PET images were Gaussian smoothed in OsiriX and smoothing was applied to raw data with a 3 x 3 matrix size and a matrix normalization value of 24. Whole lung FDG uptake was measured by first creating a whole lung region-of-interest (ROI) on the lung in the CT scan by creating a 3D growing region highlighting every voxel in the lungs between −1024 and −500 Hounsfield units. This whole lung ROI is copied and pasted to the PET scan and gaps within the ROI are filled in using a closing ROI brush tool with a structuring element radius of 4. All voxels within the lung ROI with a standard uptake value (SUV) below 1.5 are set to zero and the SUVs of the remaining voxels are summed for a total lung FDG uptake (total inflammation) value. Total FDG uptake values were normalized to back muscle FDG uptake that was measured by drawing cylinder ROIs on the back muscles adjacent to the spine at the same axial level as the carina (SUVCMR; cylinder-muscle-ratio)^38^. PET quantification values were organized in Microsoft Excel. 3D images were created using the 3D volume rendering tool on OsiriX MD.

### Luminex Isotype and FcR Binding Assay

To determine relative concentrations of antigen-specific antibody isotypes and Fc receptor binding activity, a Luminex isotype assay was performed as previously described^39^. Antigens (SARS-CoV-2 spike, RBD, S1, S2, HKU1 RBD, and OC43 RBD) were covalently coupled to Luminex microplex carboxylated bead regions (Luminex Corporation) using NHS-ester linkages with Sulfo-NHS and EDC (Thermo Fisher Scientific) according to manufacturer recommendations. Immune complexes were formed by incubating antigen-coupled beads with diluted samples. Mouse-anti-rhesus antibody detectors were then added for each antibody isotype (IgG1, IgG2, IgG3, IgG4, IgA, NIH Nonhuman Primate Reagent Resource supported by AI126683 and OD010976). Tertiary anti-mouse-IgG detector antibodies conjugated to PE were then added. FcR binding was quantified similarly by using recombinant NHP FcRs (FcγR2A-1, FcγR2A-2, FcγR3A, courtesy of Duke Protein Production Facility) conjugated to PE as secondary detectors. Flow cytometry was performed using an iQue (Intellicyt) and an S-LAB robot (PAA), and analysis was performed on IntelliCyt ForeCyt (v 8.1).

### Systems serology

To quantify antibody functionality of plasma samples, bead-based assays were used to measure antibody-dependent cellular phagocytosis (ADCP), antibody-dependent neutrophil phagocytosis (ADNP) and antibody-dependent complement deposition (ADCD), as previously described^40-43^. SARS-CoV-2 spike protein (Hexapro antigen from Erica Ollmann Saphire, La Jallo for Immunology) was coupled to fluorescent streptavidin beads (Thermo Fisher) and incubated with sera samples to allow antibody binding to occur. For ADCP, cultured human monocytes (THP-1 cell line) were incubated with immune complexes, during which phagocytosis occurred. For ADNP, primary PMBCs were isolated from whole blood using an ammonium-chloride-potassium (ACK) lysis buffer. After phagocytosis of immune complexes, neutrophils were stained with an anti-CD66b Pacific Blue detection antibody (Biolegend) prior to flow cytometry. For ADCD, lyophilized guinea pig complement (Cedarlane) was reconstituted according to manufacturer’s instructions and diluted in a gelatin veronal buffer with calcium and magnesium (Boston BioProducts). After antibody-dependent complement deposition occurred, C3 bound to immune complexes was detected with FITC-Conjugated Goat IgG Fraction to Guinea Pig Complement C3 (MP Biomedicals). For quantification of antibody-dependent NK cell activation, diluted plasma samples were incubated in Nunc MaxiSorp plates (Thermo Fisher Scientific) coated with antigen. Human NK cells were isolated the evening before using RosetteSep Human NK cell Enrichment cocktail (STEMCELL Technologies) from healthy buffy coat donors and incubated overnight with human recombinant Interleukin 15 (STEMCELL Technologies). NK cells were incubated with immune complexes, CD107a PE-Cy5 (BD), Golgi stop (BD) and Brefeldin A (BFA, Sigma-Aldrich). After incubation, cells were stained using anti-CD16 APC-Cy7 (BD), anti-CD56 PE-Cy7 (BD) and anti-CD3 Pacific Blue (BD), and then fixed (Perm A, Life Tech). Intracellular staining using anti-IFN-γ FITC (BD) and anti-MIP-1β PE (BD) was performed after permeabilizing the NK cells with Perm B (Thermo Fisher). Flow cytometry acquisition of all assays was performed using an iQue (IntelliCyt) and a S-LAB robot (PAA). For ADCP, phagocytosis events were gated on bead-positive cells. For ADNP, neutrophils were identified by gating on CD66b+ cells, phagocytosis was identified by gating on bead-positive cells. A phagocytosis score for ADCP and ADNP was calculated as (percentage of bead-positive cells) x (MFI of bead-positive cells) divided by 10,000. ADCD quantification was reported as MFI of FITC-anti-C3. For antibody-dependent NK activation, NK cells were identified by gating on CD3^-^, CD16^+^ and CD56^+^ cells. Data were reported as the percentage of cells positive for CD107a, IFN-γ, and MIP-1β.

### Statistics

The difference between any two groups at a time point was measured using a two-tailed nonparametric Mann–Whitney unpaired rank-sum test. The difference between time points within a group was measured using a Wilcoxon matched-pairs signed-rank test. All correlations were Spearman’s correlations based on ranks. All the statistical analyses were performed using GraphPad Prism v.9.0.0 or R version 3.6.1.

**Supplementary table 1.**

| **Animal ID** | **Group** | **Age at enrollment** | **Weight at enrollment (kg)** |
| --- | --- | --- | --- |
| KM71 | No vaccine | 7 | 10.3 |
| LA82 | No vaccine | 6 | 11.6 |
| LP15 | No vaccine | 4 | 4.9 |
| LD83 | No vaccine | 6 | 10 |
| A11N092 | O/W | 9 | 10.3 |
| KP93 | O/W | 7 | 10.1 |
| LR25 | O/W | 4 | 5.1 |
| LR27 | O/W | 4 | 5.8 |
| KT40 | AS03 | 7 | 10.3 |
| LA51 | AS03 | 6 | 9.5 |
| LA71 | AS03 | 6 | 10.1 |
| LR32 | AS03 | 4 | 5.8 |
| LR36 | AS03 | 4 | 6.2 |
| LB08 | AS37 | 6 | 10.9 |
| LE83 | AS37 | 6 | 11.4 |
| LB31 | AS37 | 6 | 9.5 |
| LR59 | AS37 | 4 | 6.2 |
| LR71 | AS37 | 4 | 6 |
| LB62 | CpG-Alum | 6 | 10.2 |
| LC11 | CpG-Alum | 6 | 9.7 |
| LC38 | CpG-Alum | 6 | 10.1 |
| LR89 | CpG-Alum | 4 | 5.1 |
| LT80 | CpG-Alum | 4 | 5.6 |
| LD26 | Alum | 6 | 9.1 |
| LD35 | Alum | 6 | 11 |
| LD36 | Alum | 6 | 9.6 |
| LV86 | Alum | 3 | 5 |
| MA99 | Alum | 3 | 4.6 |
| LE42 | AS03* | 6 | 11.7 |
| LH90 | AS03* | 6 | 11.2 |
| LN06 | AS03* | 4 | 5.3 |
| LN69 | AS03* | 4 | 4.6 |
| LE62 | AS03* | 6 | 8.5 |

\* Not challenged. Animals followed for durability of immune responses.

**Supplementary table 2.**

| **Fluorochrome** | **Antibody** | **Vendor** | **Catalog#** | **Clone** | **Reaction** | **Volume per reaction (μl)** |
| --- | --- | --- | --- | --- | --- | --- |
| FITC | IL-2 | Biolegend | 500304 | MQ1-17H12 | ICS | 2 |
| PerCP-eF710 | CXCR5 | Invitrogen | 46-9185-42 | MU5UBEE | Stimulation | 2.5 |
| PE | IL-4 | BioLegend | 500810 | MP4-25D2 | ICS | 1 |
| PE-CF594 | CD45RA | BD Biosciences | 565419 | 5H9 | Surface | 2 |
| PE-Cy7 | TNF- α | E-Bioscience | 25-7349-82 | Mab11 | ICS | 0.3 |
| BV421 | CD40L | Biolegend | 310824 | 24-31 | ICS | 2 |
| BV506 | TCR-γδ | Biolegend | 331220 | B1.1 | Surface | 2.5 |
| BV605 | CD4 | Biolegend | 317438 | OKT4 | Surface | 1.5 |
| BV650 | CD3 | BD Biosciences | 563916 | SP34-2 | Surface | 2.5 |
| BV711 | CCR7 | Biolegend | 353228 | G043H7 | Surface | 2 |
| BV785 | CD127 | Biolegend | 351330 | A019D5 | Surface | 2.5 |
| APC | IL-21 | BioLegend | 513008 | 3A3-N2 | ICS | 2.5 |
| A700 | IFN-γ | Biolegend | 502520 | 4S.B3 | ICS | 1 |
| APC-Cy7 | CD25 | Biolegend | 302614 | BC96 | Surface | 2 |
| BUV395 | CXCR3 | BD Biosciences | 565223 | 1C6/CXCR3 | Stimulation | 2.5 |
| BUV563 | CD8 | BD Biosciences | 612914 | RPA-T8 | Surface | 2 |
| BUV737 | CCR6 | BD Biosciences | 612780 | 11A9 | Surface | 2 |
| BUV805 | CD69 | BD Biosciences | 748763 | FN50 | Surface | 2 |

**Supplementary table 3.**

| **Animal ID** | **Group** | **Age at enrollment** | **Weight at enrollment (kg)** |
| --- | --- | --- | --- |
| LD44 | RBD-NP | 6 | 10.8 |
| LD54 | RBD-NP | 6 | 9.5 |
| MB72 | RBD-NP | 3 | 5 |
| MB84 | Soluble Hexapro | 3 | 4.5 |
| LB15 | Soluble Hexapro | 6 | 11.8 |
| LF37 | Soluble Hexapro | 6 | 11.2 |
| LI54 | Soluble Hexapro | 5 | 9.6 |
| LM34 | Soluble Hexapro | 4 | 5.1 |
| MC05 | Soluble Hexapro | 3 | 4.2 |
| LF69 | Hexapro-NP | 6 | 9.5 |
| LE43 | Hexapro-NP | 6 | 11.4 |
| LI18 | Hexapro-NP | 5 | 9.6 |
| LI22 | Hexapro-NP | 5 | 9.9 |
| LR71 | Hexapro-NP | 5 | 9.2 |
| LI34 | Hexapro-NP | 3 | 4.9 |

## Notes

### Competing Interest Statement

Derek T. O Hagan, Robbert Van Der Most and Rino Rappuoli are employees of the GSK group of companies. Robert L. Coffman and David Novack are employees of Dynavax Technologies Corporation. Harry Kleanthous is an employee of the Bill and Melinda Gates Foundation. Ching-Lin Hsieh and Jason S. McLellan are inventors on U.S. patent application no. 63/032,502 Engineered Coronavirus Spike (S) Protein and Methods of Use Thereof.

